# Brain network integration, flexibility and quasicyclicity during task and rest

**DOI:** 10.1101/2022.04.20.488888

**Authors:** Peter Fransson, Marika Strindberg

## Abstract

Previous studies have shown that a re-organization of the brain’s functional connectome expressed in terms of integration and segregation may play a pivotal role. However, it has been proven difficult to capture both processes within a single network-based framework. In this study we apply a hierarchical, spatiotemporally flexible network perspective onto fMRI data to track changes in integration and segregation in time. Our results show that network integration and segregation occur simultaneously in the brain. During task performance, global changes in synchronization between networks arise which are tied to the underlying temporal design of the experiment. We show that a hallmark property of the dynamics of the brain’s functional connectome is a presence of quasiperiodic patterns of network activation and deactivation, which during task performance becomes intertwined with the underlying temporal structure of the experimental paradigm. The proposed approach to study spatiotemporal changes in network reconfiguration during rest as well as task performance could be useful to identify aberrant network motifs in disease.

## Introduction

The use of functional magnetic resonance imaging has since its inception more than thirty years ago proven to be a powerful tool to study the relationship between behavior and activity in the brain. Early work was principally focused on anatomical localization of changes in fMRI activity using linear models. More recently, network-based models of brain activity have been introduced as a valuable compliment due to its potential to provide insights on the flow of neuronal communication between brain areas (Avena-Koenigsberger et al., 2018). The development of network-based models for fMRI experiments was largely spurred by the discovery of intrinsic and spontaneous (resting-state) networks (Biswal et al., 1995) and their subsequent use to map abnormal brain connectivity patterns related to CNS disease (Fox and Greicius, 2010). The introduction of network-based models has revitalized the question of the importance of brain network segregation versus integration and its relationship to behavior (Fair et al., 2007; Sporns et al., 2013; Deco et al., 2015; Cohen, 2017; Finc et al., 2017; Wang et al., 2021). Furthermore, the development of methods that allows for studying the time-varying properties of fMRI functional connectivity has enabled investigations of the temporal aspects of network integration and segregation (Allen et al., 2014; Preti et al., 2017; Fransson et al., 2018; Lurie et al., 2020; Kaboodvand et al., 2019). To this end, it is noteworthy that many previous time-resolved network-based models of brain segregation and integration has either implicitly (for example by underlying choices of model parameter constraints) or explicitly proposed that the brain transits through states, which to some extent, are either characterized as being integrated or segregated (e.g. Shine et al., 2015, 2016; Wig 2016; Wang et al., 2021). A more nuanced view of the integration versus segregation dictum has recently been suggested by considering brain network dynamics to be represented by PCA-driven projections of high-dimensional fMRI data into low-dimensional “manifolds” (e.g. Shine et al., 2019) and thereby achieving a continuum of brain states along the integration-segregation axis. Similarly, the hidden Markov modelling (HMM) approach have been used to divide fMRI signal intensity time-series into states and shown that transitions between states is non-random and tends to be drawn towards two sets of states that are related to higher order cognition (Vidaurre et al., 2017).

Still, there are strong reasons to assume that the processes that drive the dynamics of brain network change into either integrated or more segregated states can occur simultaneously and co-exist in the brain. It is also seems sensible to assume that spontaneously or task-driven changes in network configurations may exhibit spatial as well as temporal diversity, in the sense that co-existing networks in the brain may show a considerable variability in both spatial topology as well as modularity, i.e. integration, across time. That is, we may consider a co-existence of networks that range from being temporally fleeting to networks are more modular and integrated across time as well as networks that show traits of both types.

In a recently published study, we used resting state fMRI data to introduce a novel method that models spatiotemporally flexible networks such that networks differ both in spatial size and modularity across time (Strindberg et al 2021). In the present work we apply and extend our method to task-based fMRI data (100 subjects from the independent HCP data cohort, van Essen et al., 2012) and contrast it to the results from the analysis of resting state data. Our results that show that network integration and disintegration occur simultaneously during both tasks and rest. Further, we show that networks are repeatedly integrated and segregated, where some networks show a strong propensity for being modular, i.e. being highly integrated, whereas others are considerably more flexible and are integrated on a more intermittent basis. We show that a mapping of co-existing spatiotemporally flexible networks during task-based fMRI reveals a synchronization in time with respect to task performance. At a global level, we show that the variability in amplitude between networks causes a strong quasicyclic pattern of correlation versus anti-correlation, that in the case of task-based fMRI recordings are time-locked to the temporal structure of the paradigm. Together, our results provide insights into the relationships between network integration, spatial network flexibility, cyclicity and human behavior.

## Results

Our approach to assemble and analyze spatiotemporally flexible networks in fMRI data is founded on computing instantaneous phase synchronicity and modularity, i.e. communities (Strindberg et al., 2021). Our framework of building networks at different levels of spatial granularity, rests upon two key properties. The first being that the process of assembling networks is performed from the stand point of evaluating fluctuations in modularity in a whole-experiment perspective. Thus, we take into account the degree of membership to the same community across the entire length of the fMRI experiment rather than on a time-point by time-point basis. The second key component in our model is its combinatorial approach to the problem of how to represent both very flexible, i.e. short-lived interactions as well as temporally more stable and modular networks in the brain.

### A synthetic data example

Our approach to study network integration, flexibility and quasicyclicity in the brain is exemplified in Fig. 1. To illustrate the process of assembling together spatiotemporally flexible subnetworks based on calculations of instantaneous phase synchronization, we created a synthetic dataset that is shown in Fig. 1a. This dataset simulates 60 brain parcels that are split into three subsets, which within themselves are tightly coupled across time (120 degrees phase shift between subsets). We used instantaneous phase synchronicity analysis (IPSA) as our method of choice to compute time-resolved estimates of functional connectivity, which require the phase data to be smooth, i.e. to avoid riding waves leading to ambiguous phase information. Therefore, we used empirical mode decomposition (EMD) (Huang et al., 1998), which is a heuristic sifting algorithm that divides the signal into a finite set of oscillatory components called intrinsic mode functions (IMF) that each cover different frequency ranges in an ascending order (see also Niazy et al., 2011). For our synthetic data, the EMD algorithm generated five IMFs from which we selected the third mode function (see also IMF power spectra in Suppl. Fig. 1a). The amplitude and phase of the signal at each time point was computed by IPSA, which uses the Hilbert transform of the signal to obtain phase values at each point in time (Figs. 1b and 1c). The instantaneous phase values were then used to compute the phase coherence between all parcels at all points in time. Next, to find communities of parcels that were synchronized across time (phase coherence), we used the Louvain algorithm (Blondel et al., 2008) to estimate modularity in the data (Fig. 1d). Note that for almost all time-points, the Louvain algorithm found three (in some cases two) communities, see also Suppl. Fig. S2 that shows three distinct peaks at −1 (anti-phase), 0 (orthogonal) and +1 (in-phase).

**Figure 1.**
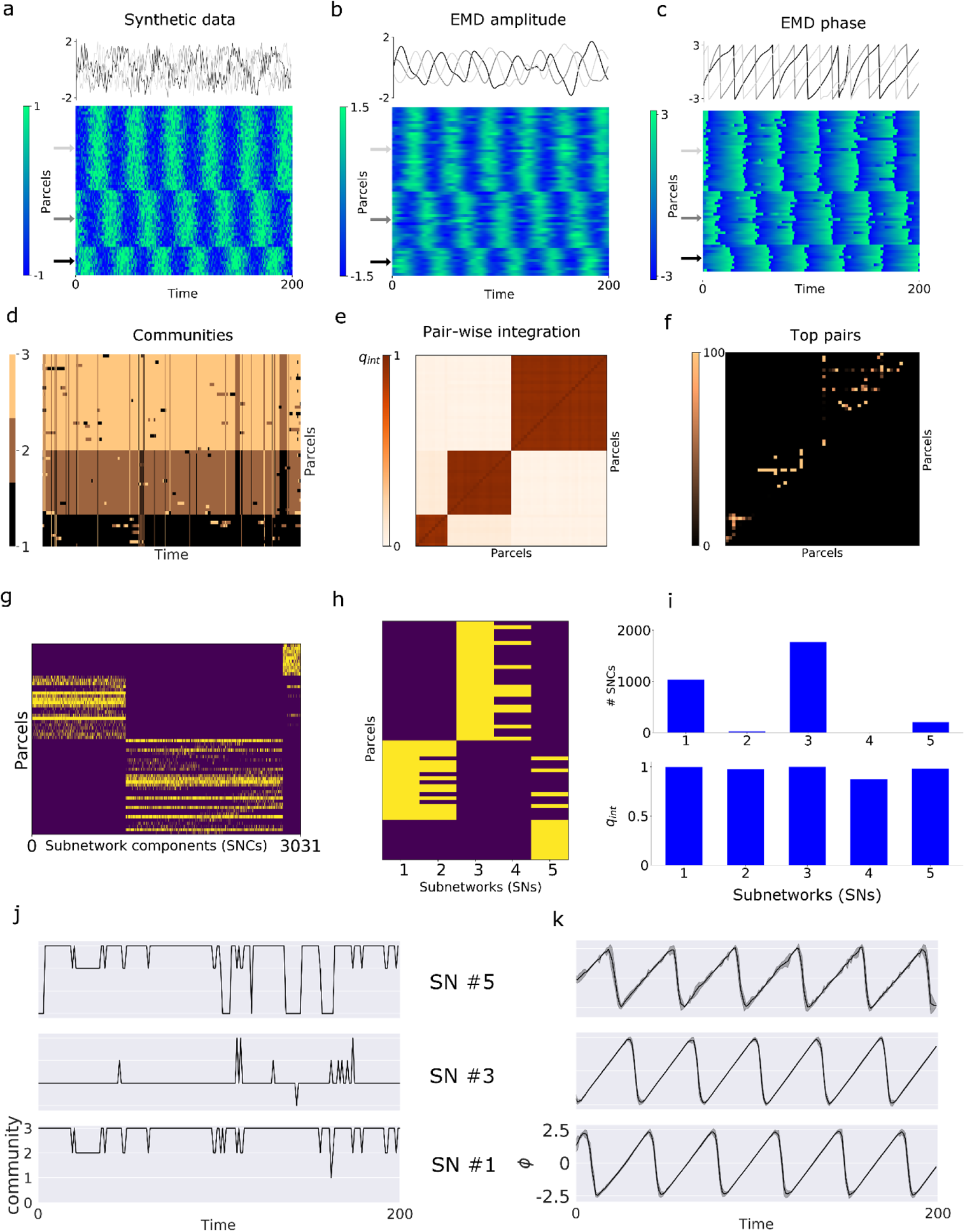
A synthetic data example that illustrate the main steps taken to compute spatiotemporally flexible network units at two levels of spatial granularity (subnetwork components (SNCs) and subnetworks (SNs)). (**a**) Input dataset (grouped into three subsets consisting of 30, 20 and 10 parcels, where each subset were generated by a sinusoidal function, separated with a phase difference of 120 degrees with added Gaussian noise, ten datasets x 400 time-points, only the first 200 time-points in one dataset shown). Data amplitude (**b**) and phase (**c**) after applying the empirical mode decomposition (EMD) algorithm. Next, community membership at each time-point for the phase data time-series was established by the Louvain algorithm (**d**) (100 runs, results from a single run shown). The key steps to compute Subnetwork components (SNCs) are portrayed in panels (**e**), (**f**) and (**g**) where we first compute the quotient (q_int_, range 0-1) defined as the duration of time each parcel is assigned to the same community relative to the total number of time-points. This was done for all possible combinations of pairs of parcels (**e**). In panel (**f**), we show the top pairs of parcels in terms of high q_int_ values (taken across multiple runs of the Louvain algorithm and each parcel is assigned to at least one pair so that all parcels are at least once represented among the top pairs). The top pairs (**f**) then acted as seeds for assembling SNCs (**g**), in which pairs of parcels were iteratively added to build SNCs (up to a size of 8 parcels) based on their q_int_ values. The final step was to group SNCs into larger integrated and spatially partly overlapping network units at a second level of granularity, which in this case resulted in five subnetworks (SNs) (**h**). Note that there is a substantial spatial overlap between SNs, in particular between SN #2 and #1 as well as between #4 and #3. In fact, the parcels included in SNs #4 and #2 are a subset of the parcels included in SNs #3 and #1, respectively. The number of SNCs included and the average q_int_ values for each SN is shown in panel (**i**). The community membership in one dataset as a function of time for the three most integrated SNs are shown in panel (**j**) (corresponding maps for individual parcels is shown in Suppl. Fig. S3). Note also that the community assignment for SNs #3, and in particular for SN #5, are at some instances in time set to zero, which implies that none of the incorporated SNCs were inherently integrated at that point in time (i.e. the SN was defined to be disintegrated at these points in time). The average phase (across the ten datasets) for the three largest SNs are shown in panel (**k**) (see also Supplementary Methods and Suppl. Fig S3).

We have previously shown that a positive phase relationship between two brain areas correspond to an assignment to the same community, whereas assignment to different communities implies negative (or orthogonal) phase synchronicity (Strindberg et al., 2021). Thus, a frequent assignment of parcels to the same community is therefore coupled to a positive phase relationship across time whereas segregated parcels were assumed to have an orthogonal or negative phase coherence. Based on this relationship, we introduce the key parameter in our model, q_int_ which is the quotient between the number of time-points a pair of parcels belong to the same community divided by the total number of time-points for the entire fMRI scan (Strindberg et al., 2021). At a later stage, we will re-use the same concept to assemble network units at two different levels of spatial granularity, subnetwork components (SNCs) and subnetworks (SNs). Thus, q_int_ is a parameter that measures the temporal coupling strength between parcels and takes on values between 0 and 1, where 0 implies no time-points of assignment to the same community in any subject and 1 means an assignment to the same community for all time-points throughout the scan in all subjects. In terms of networks, a high q_int_ value implies a high degree of (within)-integration whereas a low value signifies a high degree of disintegration (i.e. parcels are assigned to different communities). The matrix displayed in Fig. 1e. shows pair-wise measures of q_int_ values for all parcels in our synthetic data example. As shown in Fig. 1e., we observe high q_int_ values for pairs of parcels belonging to the same subset and low q_int_ values for pairs situated in different subsets. Fig. 1f shows the top ranked pairs of parcels with the highest q_int_, selected so that each parcel is represented at least once (taken across 100 runs of the Louvain algorithm).

The top-pairs shown in Fig. 1f were then used as seeds in a combinatorial approach to assemble subnetwork components (SNCs), which stretches beyond of pairs of parcels and therefore constitute an intermediate step in terms of taking into account different spatial levels of network granularity in our model. The rationale for using a combinatorial approach was make sure that we could adequately capture both modular and more flexible coupling mechanisms among all parcels (Strindberg et al., 2021). Modularity here refers to a preference of the phase of a parcel to be synchronized with the phase of the same set of parcels throughout the scan. On the other hand, flexibility refers to the tendency of parcels to preferentially tend to synchronize with different sets of parcels during the scan. Thus, pairs with high q_int_ values tend to be modular, while pairs with low values tend to be more flexible across time. It is very likely that parcels vary in their tendency towards being modular or flexible towards other parcels and it is therefore important that the strongest phase synchronizations from the point of view of each individual parcel is represented from the start in the process of assembling SNCs and, at the second level of granularity, the merge of SNCs into SNs (see below). By preserving the strongest phase synchronization for each parcel, we wanted to avoid the scenario that modularity dominates over flexibility by design. Therefore, SNCs were assembled in an iterative, stepwise fashion, starting with the top pairs shown in Fig. 1f, so that all parcels in the parcellation scheme are represented at each level. The size of the SNCs were iteratively increased by a factor of 2 until the final size of 8 parcels was reached (Fig. 1g) (see also Supplementary Methods). In our synthetic data example, this resulted in 3031 unique, but partly overlapping SNCs, where each SNC has its own subject-specific time-course of community membership. The time-courses of community membership were used to compute q_int_ values that represents the average degree of temporal coupling strength for each SNC. As shown in Fig. 1g, the large majority of SNCs represent phase synchronization within each subset of parcels and they tend to overlap to a large extent with each other (see also Fig. 1d), although a smaller number of SNCs spanned parcels residing in different subsets. Note that SNCs can internally be in a state of integration (all parcels assigned to the same community) or disintegration (parcels lack internal phase coherence and are scattered across communities). Moreover, SNCs can be said to be segregated vis-à-vis other SNCs by the fact that their corresponding phase (averaged across parcels) is either orthogonal (phase equal to zero) or negatively correlated (anti-phase). A pictorial description is given in Suppl. Fig. S4 in Strindberg et al., 2021).

At the second spatial granularity level, we grouped together SNCs into subnetworks (SN) that were allowed to vary in spatial size (Fig. 1h). This grouping was based on the criteria that all SNCs assembled into the same SN were guaranteed to be assigned to the same community (Fig. 1d) at every time-point of integration. Thus, each SN had a subject specific time-course of community assignment and a time-resolved spatial configuration that resulted from combining the parcels of all the SNCs into a common time-resolved representation of a SN. A SN was considered to be integrated at a given point in time if at least one of its constituting SNCs were integrated (i.e. all parcels in the SNC were assigned to same community). Consequently, if all SNCs in a given SN at a given point in time was classified as being disintegrated, then the SN was also considered to be disintegrated at that particular point in time. The fact that the spatial configuration of SNs varies over time implies that some parcels will often or always be part of a specific SN while other parcels will more rarely make a contribution. This kind of probabilistic and combinatorial representation could theoretically lead to a scenario where all parcels take part in all SNs or ultimately create one large undifferentiated SN. Therefore, in order to penalize these undesired scenarios, a criterion of internal coherence of the SNs was used to guide the process of merging SNCs into SNs. Internal coherence meant that none of the SNCs that together formed an SN could at any point in time be segregated vis-á-vis each other (i.e. showing anti-phase or orthogonality in phase which would result in integration simultaneously in different communities). For further details of how SNs are assembled from SNCs, see Supplementary Methods.

Turning back to our synthetic data example, we observe that the assembly of SNs built from SNCs yielded five SNs (Figs. 1h and 1i) of which three (SN #1, #3 and SN #5) are very closely associated with the three subsets in the input data shown in Fig. 1a. The three aforementioned SNs together incorporates 99.2 percent of all available SNCs and they are highly integrated throughout the scan (mean q_int_ values = 0.999, 0.999 and 0.981), i.e. at least one of the incorporated SNCs is integrated at each time-point. This fact is also reflected in the community assignment time-courses shown in Fig. 1j for one dataset, where a community value of 0 means non-integration (disintegration), i.e. none of SNCs included in the corresponding SN were internally integrated (i.e. all of the 8 parcels in any SNC were assigned to the same community) at that point in time. Finally, the mean phase for all parcels included in each of the three dominating SNs are shown in Fig. 1k, which are closely matched with the input data shown in Fig. 1a-c. It is important to point out that, within the modelling framework presented here, integration and segregation are not mutually exclusive events in the temporal domain. Rather, at any given point in time, two different ensembles of parcels can each be integrated into separate communities, but at the same time segregated relative to each other. This implies that we can model integration as well as segregation that take place simultaneously in time between different networks. To be clear, when segregation happens within the same network it is referred to as disintegration when means that the network is not in fact assembled/active due to a lack positive internal phase coherence. Thus, integration and disintegration are mutually exclusive in the temporal domain. A more detailed picture of the spatiotemporally flexible properties of the subnetworks is provided in Suppl. Fig. S3 that shows how the distribution of participating parcels in all five subnetworks changes as a function of time as well as their community membership.

### Assembling spatiotemporally flexible subnetworks

Equipped with the insights gained from the synthetic data example, we proceed by presenting the preparatory results in the case of the working memory HCP dataset shown in Fig 2. The EMD sifting algorithm resulted in four intrinsic modes of which we selected the third mode (IMF3, covered the 0.01 – 0.1 Hz frequency range, see also Suppl. Fig. 1b). Similar to the synthetic data example, the instantaneous phase synchronization analysis produced three peaks of phase synchronization values at +1, 0 and −1 (Suppl. Fig. 2b) which resulted in that the Louvain algorithm divided the phase synchronization values into three communities at most time-points (Fig. 2d). The pair-wise distribution of q_int_ values based on assignment to communities for all possible pairs of parcels is shown in Fig. 2e (q_int_ range: 0 – 0.63) and the top pairs (244 pairs) across 100 runs of the Louvain algorithm are shown in Fig. 2f. Analogously to the synthetic data example, from the top pairs we first iteratively built SNCs (30506 SNCs, size = 8 parcels) (Fig. 2g), and then merged the SNCs into 375 SNs (Fig. 2h). The number of SNCs that were assembled into each SN is shown the upper graph in Fig. 2i and the corresponding mean (across subjects) q_int_ value for each SN is shown in the lower graph. From the results shown in Fig. 2i we see that a fairly small number of SNs encapsulates the majority of SNCs, while the large majority of SNs contain very few or only a single SNC.

**Figure 2.**
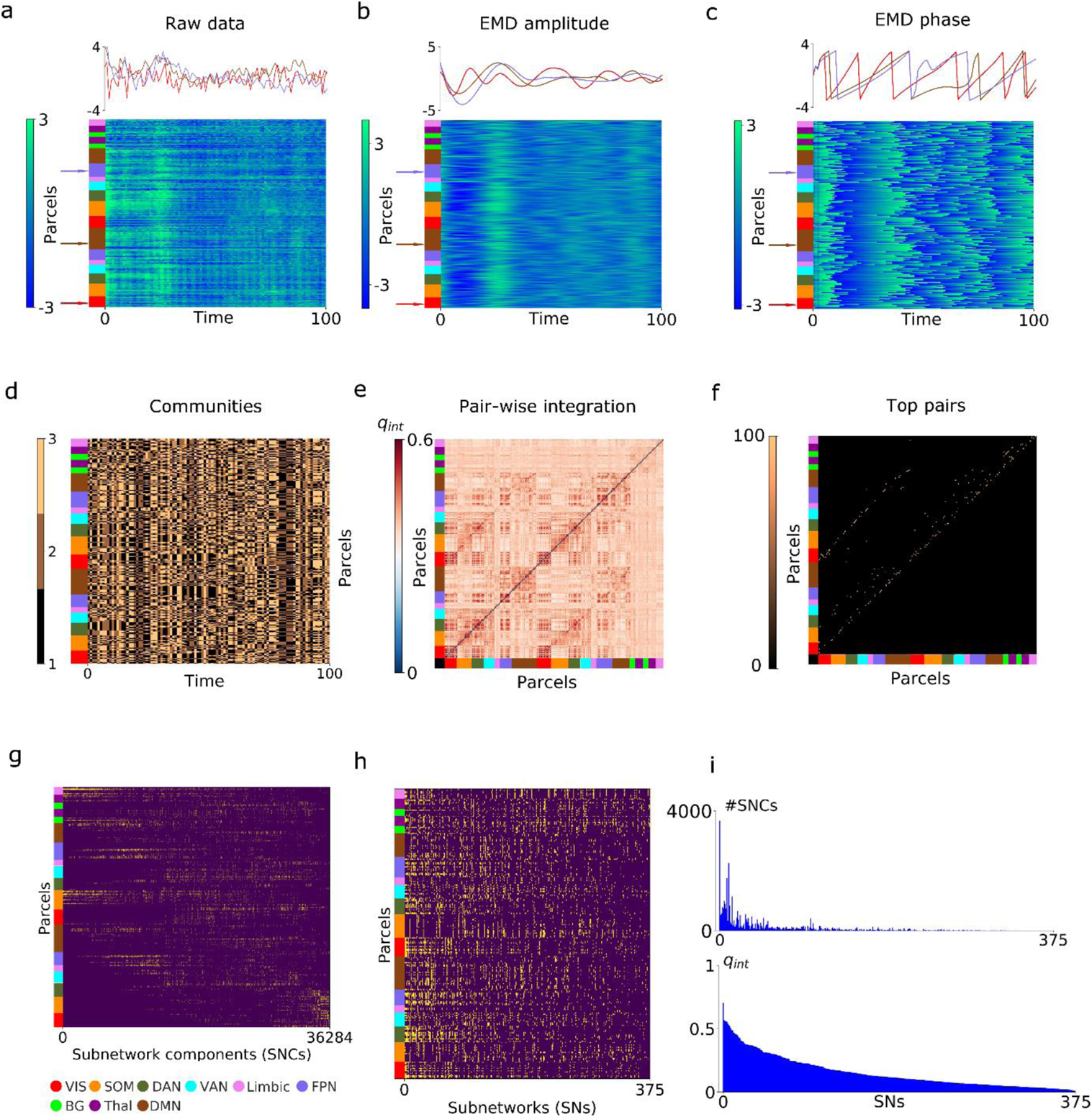
Assembly of subnetwork components (SNCs) and subnetworks (SNs) for the working memory dataset. The layout here follows closely the synthetic data example shown in Fig. 1. (**a**) Raw data for the first 100 (of 405) data-points for all parcels in one subject followed by the amplitude (**b**) and phase (**c**) time-series after empirical mode decomposition. (**d**) Division of parcels into communities by means of the Louvain algorithm (1 run out of 100 shown). The pair-wise integration matrix (**e**) shows that the degree of integration for the most pair was q_int_ = 0.6, which implies an assignment of both parcels to the same community during sixty percent of the total duration of the working memory fMRI task experiment. Top pairs of integration were largely found for parcels located within same the canonical resting-state networks, i.e. along the diagonal in panel (**f**). The iterative process of assembling pairs of parcels into subnetwork components resulted in 30506 unique, but partially overlapping subnetwork components (SNC). The final step of assembling SNCs into larger network units that are integrated but also flexible yielded 375 subnetworks (SNs) (panel (**h**), sorted by their q_int_ values in descending order). The histograms shown in panel (**i**) shows that largest SN (3665 SNCs assembled into a single SN) also had the largest qint value (q_int_ = 0.70, i.e. this SN was on average integrated during 70 percent of the total duration of the working memory experiment). Note also that only relatively few SNs are assembled from many SNCs and that the large majority of SNs are assembled from just a few SNCs, or in the extreme case, from a single SNC.

The iterative assembling of SNCs and subsequent build of SNs for the motor and rest fMRI data sets yielded 28559 SNCs, 282 SNs (motor) and 26566 SNCs, 238 SNs (rest) (data not shown).

The spatial topology and the relative contribution from each of the nine canonical RSN and the mean (across subjects) phase time-course for the top five SNs (i.e. the most integrated subnetworks across time and therefore have the highest q_int_ values) are shown in Fig. 3. For the n-back working memory task (Fig. 3a), all top five SNs have very similar spatial topology that included visual, medial premotor cortex, lateral posterior parietal, dorsolateral prefrontal cortex and thalamus. All are regions that previously have been consistently associated with n-back tasks (Owen et al., 2005). Unsurprisingly, the top five SNs displayed very similar temporal phase profiles marked by positive phase peaks during the intermittent epochs of rest. While the top five SNs in the working memory task are spatially very similar (Fig. 3a), they are not identical. By this we mean that they could briefly segregate (i.e. have phase profiles that tend towards orthogonality or anti-phase vis-á-vi each other) at time points when they don’t share any common parcels.

**Figure 3.**
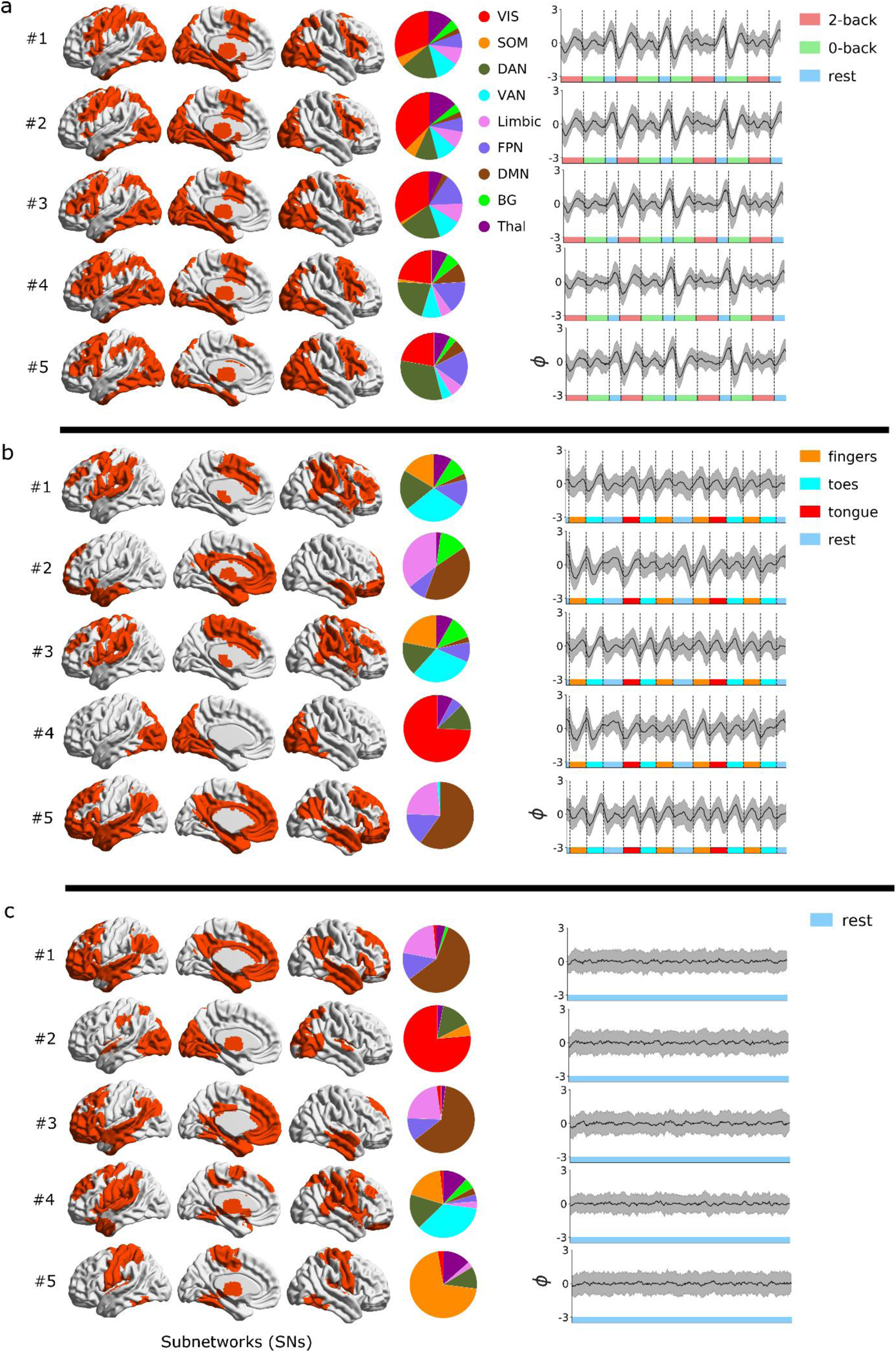
Spatiotemporally flexible subnetworks (SNs) assembled during working memory (**a**), motor (**b**) and rest (**c**). The figure shows the top five most integrated SNs (i.e. the SNs had the largest mean q_int_ values). The pie charts in the middle column show the relative contribution (dice coefficient) from the nine canonical resting-state networks to each SN. The plots to the right show the average (across subjects) phase time-course for each SN. During the working memory task (**a**), the top five SNs included foremost parcels in the visual, fronto-parietal, dorsal/ventral attention, basal ganglia and thalamic networks (mean q_int_ = 0.70, 0.57, 0.56, 0.56, 0.56; number of SNCs assembled: 3665, 522, 587, 786 and 745). The top five SNs assembled during the working memory task are spatially congruent and hence also display similar phase signal intensity time-courses. During the motor task (**b**), parcels from the somatomotor network are included in two of the five top SNs (#1 and #3) but we also find SNs that are spatially similar to canonical RSNs such as the default mode network (SNs #2 and #5) and the visual network (SN #4) (mean q_int_ = 0.62, 0.57, 0.55, 0.55, 0.5; number of SNCs assembled: 1602, 1250, 1135, 894, 1763). The phase time-courses during the motor task are rather similar across SNs, but to a lesser degree compared to the working memory experiment. For rest (**c**), the top five SNs largely conform to the canonical resting-state networks, that is default mode network (SN #1 and #3), visual network (SN #2), ventral attention network (SN #4) and somatomotor network (SN #5) (mean q_int_ = 0.62, 0.57, 0.55, 0.55, 0.51; number of assembled SNCs: 2588, 825, 1360, 2443, 992). Due to the fact that rest is not conditioned to external stimuli, the phase data time-series cancel each other out when averaged across subjects.

In contrast, the topology for the top five most integrated SNs during the motor task were spatially heterogeneous (Fig. 3b). Both SN #1 and #3 were largely assembled from parcels belonging to the somatomotor, dorsal and ventral attention networks and the basal ganglia. In contrast, SN #2 and #5 included a large contribution of parcels situated in the default mode network. The majority of parcels assembled together in SN #4 was located in the visual network. In the case of rest (Fig. 3c), four of most integrated SNs (SNs #1, 2, 3, and 5) to a large extent resembled canonical RSN (SN #1 – DMN, SN #2 – visual network, SN #3 – anterior DMN and SN #4 – somatomotor network). The remaining top SN (SN #4) formed during rest was assembled by parcels located in the ventral and dorsal attention networks, somatomotor network, and to a lesser degree, thalamus and basal ganglia. Because the rest experiment is not time-locked to any external stimuli, the mean phase time-courses (averaged across subjects) are close to zero for all data-points. For an animated example of fluctuations phase in both space and time for SNs across the entire experiments in a single subject, see Supplementary Movie M1 (SN #1, working memory), M2 (SN #1, motor) and M3 (SN #1, rest). Importantly, the movies clearly show that the number of participating parcels in a given SN varies across time. Thus, the spatial variability in subnetwork activity is dependent on the number of incorporated SNCs that are integrated into the same community at each given point in time (i.e. compare with toy data results in Suppl. Fig. S3).

To grasp the full extent of the hierarchical assembly of spatiotemporally flexible subnetworks, it is of interest to compare the properties of the most integrated (Fig. 3) to the least integrated SNs (see Suppl. Fig. 4). Whereas the top SNs are assembled from hundreds and even in some cases thousands of SNCs (see also Fig. 2i), the SNs at the bottom consist in many cases of only a single or, in one case, three SNCs. Top integrated SNs are on average integrated for more than 50 percent of the scanning time (even up to 70 percent for the working memory experiment), while the bottom SNs are integrated during less than 3 percent of the time. See also Suppl. Fig. S5 for an example of the temporal evolution of community membership, i.e. integrated versus disintegrated points in time for the top five (Suppl. Figs. S5a-c) and the five least integrated (Suppl. Figs. S5d-f) SNs in a single subject. These results suggest that our parsimonious approach to assemble spatiotemporally flexible subnetworks based on computing phase synchronization between pairs of parcels and then using a combinatorial scheme to bin them together at increasingly coarser spatial scales, is able to capture key features of subnetwork integration and disintegration and that they to a varying degree show a propensity be to synchronized during task fMRI experiments.

### How frequently and for how long do subnetworks stay integrated in time?

The results exemplified in Suppl. Fig. S5 suggests that subnetworks oscillate back and forth from being disintegrated (i.e. none of the assembled SNCs are internally integrated) to being integrated (at least one of assembled SNC is internally integrated). How often do subnetworks integrate and how long do they stay integrated before they oscillate back to a period of disintegration? This question is addressed in Fig. 4. that plots (average across subjects) the number of instances of integration (x-axis) versus duration of integration periods for all subnetworks (y-axis). Interestingly, the mean duration of integration periods for subnetworks is very similar across tasks (memory: 3.8 TRs, motor: 3.8 TRs and rest: 3.6 TRs). We note that for the memory task, the top most integrated subnetwork (SN #1) stands out with periods of integration that on average lasts for 9.3 TRs (Fig. 4a). Although the results shown in Fig. 4. seems to suggest that the average (across subjects) number of instances of integration is smaller for the motor task (10.6) than working memory (14.4) and rest (16.7). However, this is mainly an effect of the shorter experiment time span for the motor (284 TRs) compared to rest (405 TRs, only the first 405 of 1200 data-points were used in the analysis) and working memory (405 TRs).

**Figure 4.**
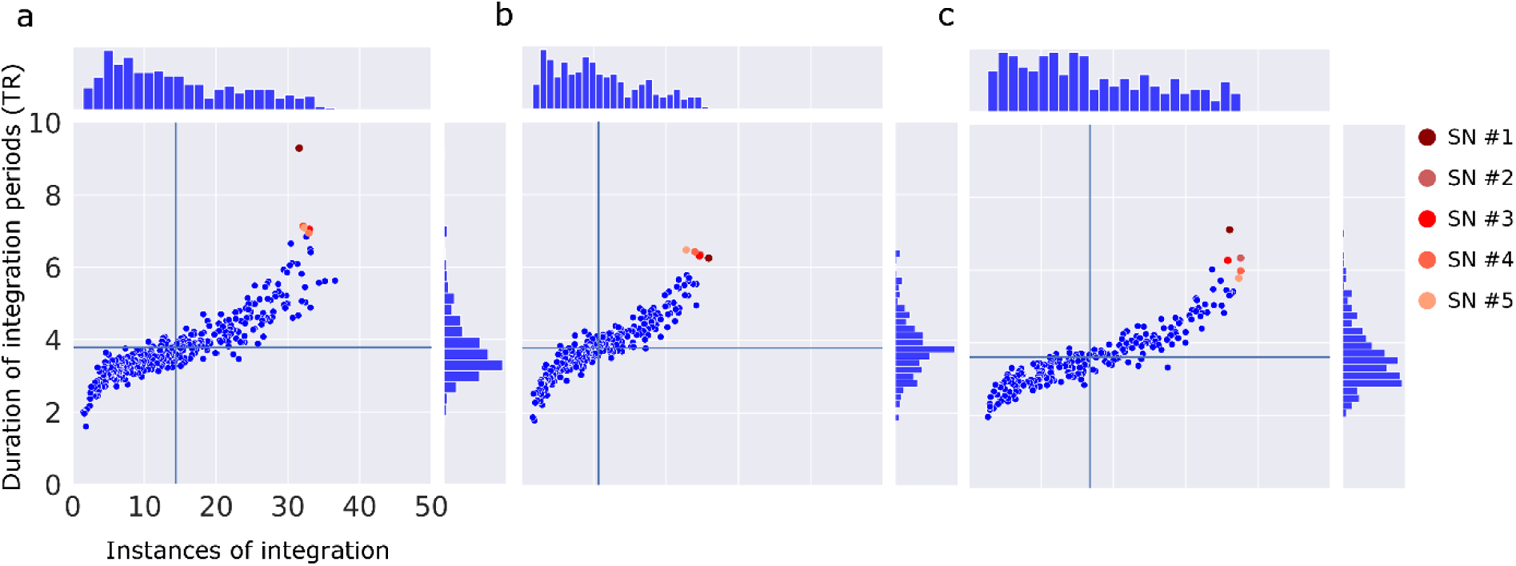
Temporal characteristics of subnetwork (SN) integration during working (**a**), motor (**b**) and rest (**c**). The x-axis shows the average (across subjects) number of instances of integration for each SN. The y-axis shows average duration (across subjects) of instances of integration. The top five SNs are marked by color. Solid lines mark the average (across SNs and subjects) number of instances of integration and duration of integration. SN #1 for the working memory task showed a marked prolongation of the average degree of integration (9.3 TRs = 6.7 seconds). For all tasks, the relationship between number of integrations and duration of integration is positive.

Another interesting question is whether the temporal pattern of integration versus disintegration (as exemplified in a single subject in Suppl. Fig. S5) is associated with the timing and execution of different tasks in the motor and working memory experiments. Suppl. Fig. S6 shows for each subnetwork and time-point the degree of integration averaged across subjects together with plots of their q_int_ values and number of subnetwork components (SNCs) included. A key observation from these results is the marked increase in subnetwork integration during the intermittent rest epochs for the working memory experiment (Suppl. Fig. S6a). Interestingly, the increased level of subnetwork integration is maintained during the directly following 0/2-back task epoch. This pattern is most prominent in the subnetworks that display the overall highest level of integration (i.e. large q_int_ values). The corresponding results for the motor task does not show similar signs of temporal patterns of integration among subnetworks (Suppl. Fig. S6b). As expected for the rest experiment, we observe an absence of any temporal regularity of subnetwork integration across the entire fMRI scan (Suppl. Fig. S6c).

### Modularity and flexibility at the level of individual parcels

To relate properties of subnetworks to lower levels of granularity, i.e. subnetwork components (SNCs) and individual parcels, we computed parcellated estimates of spatial diversity and temporal participation. We defined spatial diversity as the proportion of the total number of parcels (from a total of 236 parcels) that each individual parcel shared an SNC with. Theoretically, spatial diversity lies within the interval [(8-1)/236, 235/236] = [0.03, 0.99], where the lower limit is reached when a parcel participates in only one SNC (size of SNCs = 8 parcels), and the higher limit is reached when a parcel shares SNCs with all other parcels. Further, we defined temporal participation (or equivalently, modularity) as the relative frequency of participation of a parcel in any SNC across the entire experiment (i.e. the amount of time the parcel is represented within our ensemble of SNCs). The theoretical interval for the participation parameter is [1 / (405 * 100), 1] = [0.000025, 1], where the upper limit is reached when an area is represented in any SNC at all time-points in all of 100 subjects included in the study. The criteria for the lower limit is reached when a given parcel is integrated into any SNC only once in a single subject.

The degree of spatial diversity and temporal participation for individual parcels (averaged across subjects) are presented in the left column in Fig. 5, where temporal participation (x-axis) is plotted against spatial diversity (y-axis). The corresponding values are superimposed onto the cortical surface shown in the right column. Each spatial diversity/temporal participation graph is divided into four quadrants that are labeled hyper-flexible, flexible, flexible-modular and modular. This division is done for conceptual clarity but should be understood in terms of gradients. Parcels with a strong modular tendency score high on temporal participation but low on spatial diversity (lower right quadrant). This is because they are represented by a relatively small set of spatially homogenous SNCs that frequently integrate. In contrast, parcels with high flexibility typically are represented by a larger set of more diverse SNCs that tend to integrate briefly (upper left corner). Some parcels are represented by both types of SNCs (upper right corner). Others are represented by only a small number of SNCs each with low q_int_ (lower left corner). In the latter case the SNCs could in principle be equally diverse as those in the upper left corner but due to the smaller number SNCs the parcels will score lower on spatial diversity.

**Figure 5.**
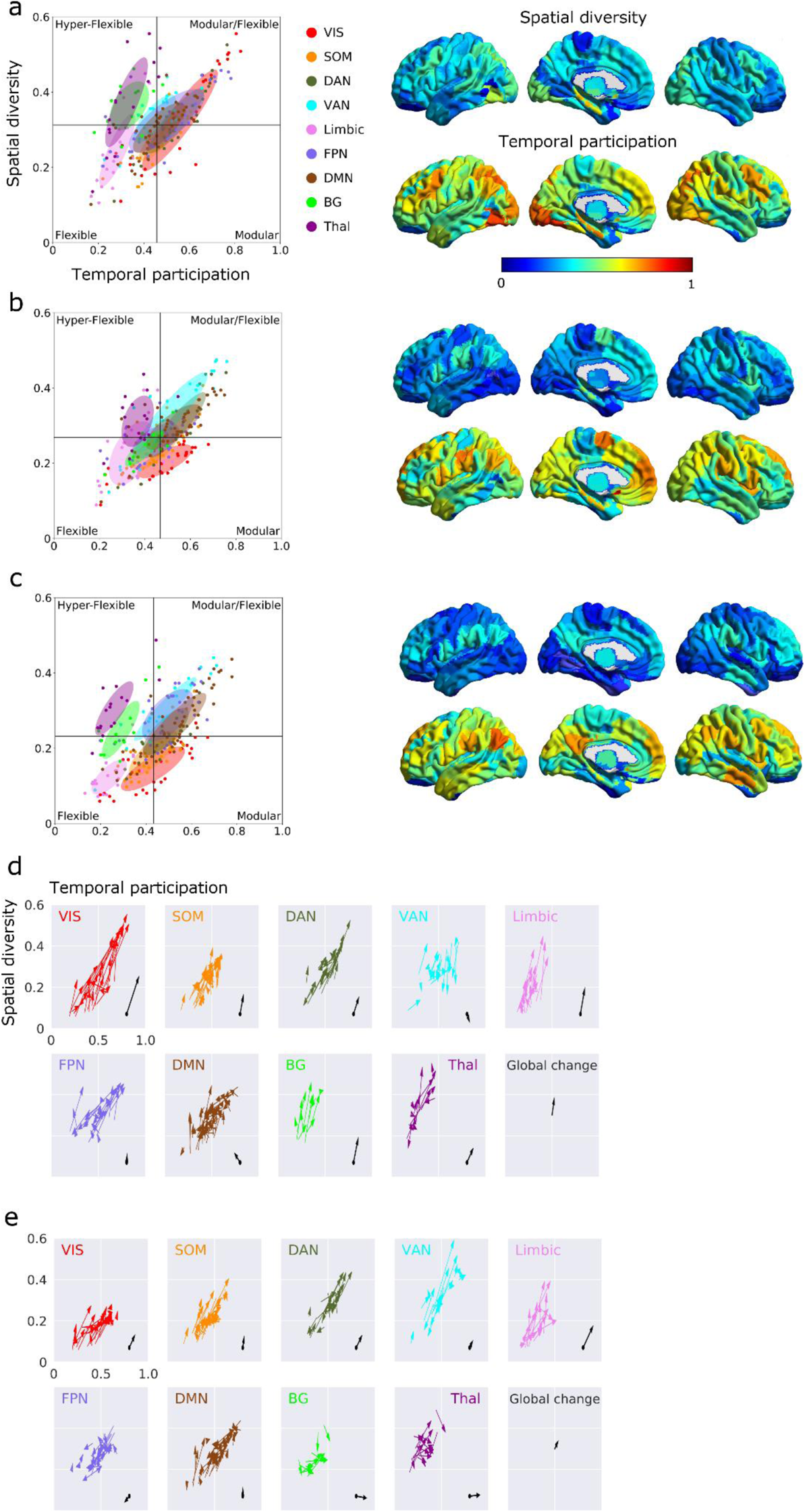
Spatial diversity and temporal participation calculated at the level of individual parcels for working memory (**a**), motor (**b**) and rest (**c**). Solid vertical and horizontal lines marks the mean values for spatial diversity and temporal participation. Individual parcels can be denoted as being either hyper-flexible, flexible, modular-flexible or modular (see text). The difference between working memory and rest in terms of spatial diversity and temporal participation at the level of individual areas (divided up into canonical RSNs) are shown in panel (**d**). Analogously, the difference between motor task and rest is shown in panel (**e**). Each ellipse in panels **a**-**c** represent all parcels belonging to a specific canonical RSN and is centered at its mean value, where the width and height corresponds to two standard deviations of the mean. The tilts of the ellipses are set by their co-variance. The tail of each arrow in (**d-e**) mark the degree of spatial diversity and temporal participation measured during rest. The head of the arrow marks the degree of spatial diversity and temporal participation during working memory (**d**) and motor task (**e**). The average change of spatial diversity and temporal participation within each canonical RSN are marked with a black arrow in the lower left corner.

The maps shown in Fig. 5a-c shows that spatial diversity and temporal participation are heterogeneously distributed between as well as within canonical RSNs. During rest (Fig. 5c), many parcels in the default mode, fronto-parietal and ventral attention networks falls into the flexible-modular category. That is, they are incorporated into SNCs that bind together a higher diversity of parcels (high spatial diversity) while at the same time being integrated for longer time-periods (high degree of temporal participation) than average (solid lines in the left graphs in Fig. 5a-c). In contrast, almost all parcels in the thalamic and many parcels in the basal ganglia network are categorized as hyper-flexible, implying similar levels of spatial diversity as the flexible-modular areas but a lower frequency of integration across time (decreased temporal participation). All parcels in the limbic network are categorized as flexible. This implies that they are represented by a smaller number of SNCs that all have low q_int_ values. The low q_int_ values is a reflection of the tendency of the limbic areas to continuously integrate in novel constellations. Parts of the visual and somatomotor networks are characterized as being modular, which suggests a high degree of temporal participation together with a low spatial diversity (i.e. they preferentially repeatedly integrate with the same small set of parcels).

To facilitate a comparison between experiments, the subpanels in Figs. 5d and 5e show the difference in spatial diversity and temporal participation between working memory and rest (Fig. 5d) as well as between motor and rest (Fig. 5e). Interestingly, during task performance, and in particular for the memory task, the degree of spatial diversity is increased for many parcels located in the visual, dorsal attention, limbic (including the hippocampus) and the basal ganglia network (Fig. 5d). This finding implies that tasks, and in particular for the working memory task, impose an increased spatial flexibility by participating in new subnetwork components in comparison to rest. The change in temporal participation due to task performance was overall smaller, with the exception of the visual, and to a smaller extent, the basal ganglia, limbic and thalamic networks, which showed an overall increase in temporal participation. That is, parcels in these networks were integrated into a SNC for substantially longer time periods during the memory task compared to rest. Additionally, our results suggest that parcels located within the visual, dorsal attention, limbic and the basal ganglia shares SNCs with a larger number of other parcels during task performance compared to rest. Of note, changes in spatial diversity and temporal participation as a function of task are considerable at the level of individual parcels, which highlights the fact that task induced changes in network flexibility and modularity is not dependent on the anatomical constraints imposed by the canonical resting-state networks. For both tasks and rest, we found (across all parcels) a positive correlation between the degree of parcel integration and spatial diversity (working memory: r = 0.61, motor: r=0.60 and rest r = 0.61; Spearman rank correlation, p<0.0001). The positive relationship between temporal participation and spatial flexibility implies that our model captures the full spectrum of interactions better for modular than for the more flexible parcels.

### Global quasicyclicity of subnetwork activation and deactivation across time

Summaries of global patterns of intrinsic co-activity are often provided in the format of a N x N matrix, where N is the number of parcels. This presentation gives an overall view of correlations that exist in the two-dimensional N by T data matrix, where T is the number of time-points. However, we can in a similar manner detect changes in global brain subnetwork configuration by computing the correlation between the amplitudes of BOLD signals across subnetworks at different points in time. So, rather than computing pair-wise correlation across time, we instead computed the correlation coefficient between subnetwork amplitudes for all combinations of time-points. This results in a T by T matrix (i.e. a recurrence matrix) where a given matrix element represents the degree of subnetwork amplitude correlation or anti-correlation (after signal demeaning) for any given pair of time-points. Of note, the recurrence matrix elements close to or at the diagonal will be strongly positive due to the auto-correlation of the BOLD response. The recurrence matrix provides a simple means to estimate the combined effect of activation and de-activation of all subnetworks at any points in time.

Recurrence matrices for all experiments at the level of SNs computed at the single subject level and then averaged across subjects are shown in Figs. 6a-c (of note, it is also possible compute recurrence matrices at the level of parcels, see Suppl. Fig. S7). In particular, the amplitude recurrence matrix for the working memory task (Fig. 6a) shows a strong and quasicyclical pattern of globally repeated patterns of correlation and anti-correlation in signal amplitude between subnetworks during the entire experiment. Quasicyclicity in amplitude activation/deactivation across subnetworks is also present during the motor task (Fig. 6b), albeit weaker in strength. In the case of rest (Fig. 6c), there are no discernable patterns of quasicyclicity due to the fact that rest is not constrained by instructions given to subjects or to otherwise externally given stimuli. However, at the level of individual subjects, quasicyclicity is present at rest, see Suppl. Fig. S7. We also note that in between periods of correlation and anticorrelation, there exist short, intermittent lapses of orthogonality, i.e. the recurrence matrix values take on values close to zero.

**Figure 6.**
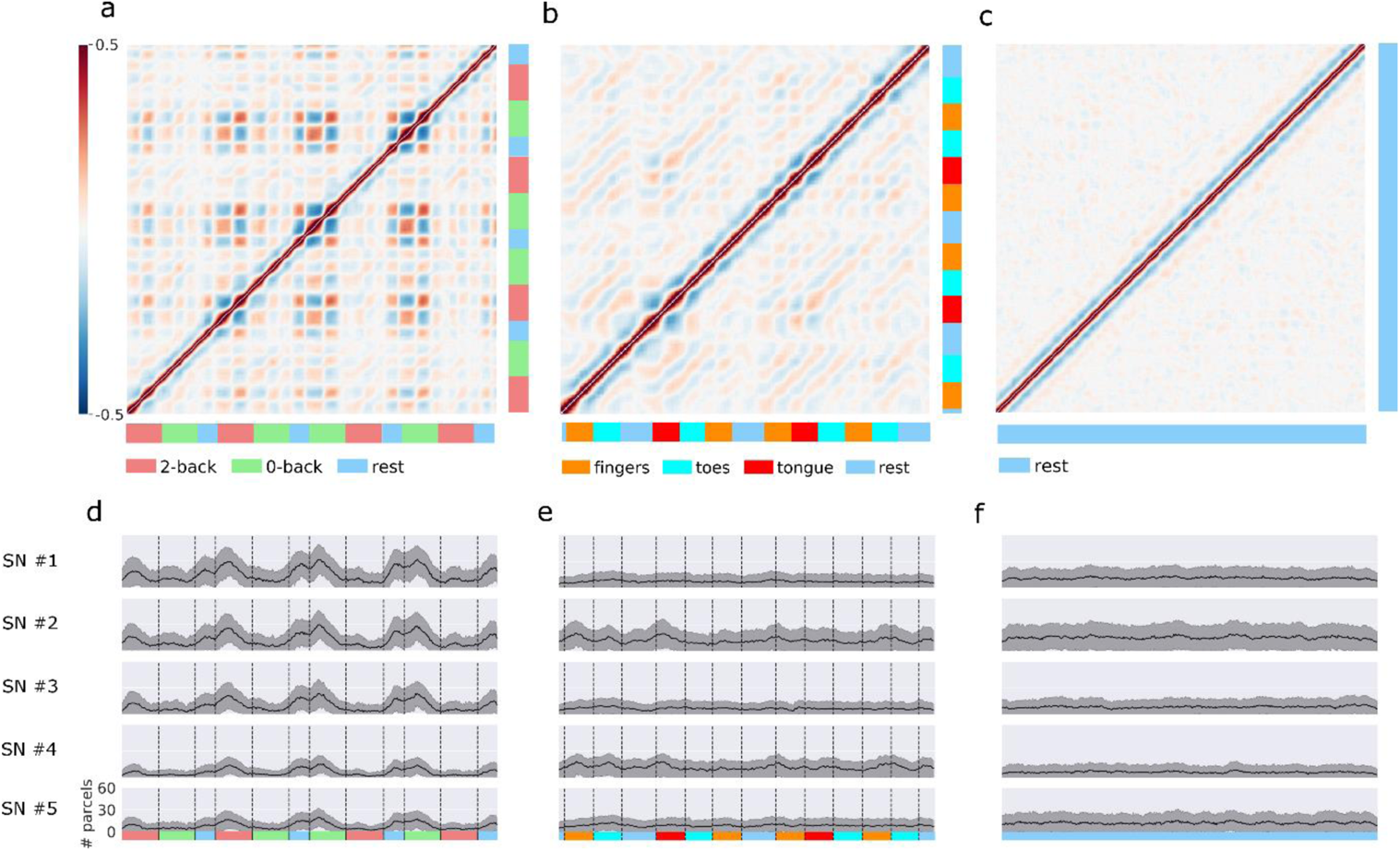
Recurrence matrices that shows the degree of correlation in amplitude activation and de-activation across subnetworks (SNs) during working memory (**a**), motor (**b**) and rest (**c**) at different points in time. Panels **d-f** shows the mean (across subjects) number of parcels participating in the top five most integrated subnetworks as a function of time. A strong pattern of quasicyclic pattern of amplitude correlation and anti-correlation between all subnetworks are seen for the working memory experiment (**a**), and to a lesser extent during the motor experiment (**b**). Time-locked quasicyclicity of subnetwork amplitude configurations is absent for the average rest recurrence matrix (**c**) (but see also Suppl. Fig. 7 for results obtained in single subjects). Note also that the strongest patterns of anti-correlation and correlations for the working memory task coincide in time with large increases in the number of participating parcels in the top five most integrated subnetworks.

The recurring pattern of correlation and anticorrelation during the working memory task, starting during the intermittent rest epochs and then continue throughout the subsequent 0/2-back epoch stands out from the patterns observed during motor task and rest (Fig. 6a). To gather insight on what might be a contributing factor to this pattern, we plotted the mean (across subjects) number of participating and integrated parcels for the top five SNs (see also Figs. 3a and 4a) as a function of time (Figs. 6d-f). In comparison to motor and rest, the top five integrated SNs for the memory task show considerably elevated numbers of participating parcels, typically showing an average increase of 200 percent or more (i.e. from 10 up to 30 parcels or more) during periods of increased correlation and anti-correlation (Figs. 6a-c).

### Subnetwork integration and task performance during the working memory experiment

The large variability in global activation/deactivation patterns among subnetworks (Fig. 6a) and fluctuations of the number of participating parcels during the time-course of the working memory experiment (Fig. 6d) warranted us to perform an investigation of putative links between subject performance (i.e. number of wrong responses given during the 0- and 2-back epochs) and the number of disintegrated time-points for the most integrated subnetwork (SN #1). Our result that subnetwork quasicyclicity is associated with the temporal order of task and rest epochs in the working memory experiment (Figs. 6a,d), prompted us to divided up responses not only with respect to task difficulty (0-back versus 2-back) but also with respect to their temporal order within the paradigm. The results are summarized in Fig. 7, which perhaps not surprisingly, show a significant increase in wrong answers given by the subjects during epochs of 2-back compared to 0-back tasks (Fig. 7a). Interestingly, this finding was mirrored in terms of a significantly higher number of disintegrated time-points for the top subnetwork (SN #1) during 2-back compared to 0-back tasks epochs (Fig. 7c). Moreover, the amount of non-integrated time-points for the top SN was significantly higher during 2^nd^ epochs compared to 1^st^ epochs (Fig. 7b; see also Fig. 7c for a definition of 1^st^ and 2^nd^ epochs). However, there was no significant difference in the number of wrong answers provided by the subjects given during 1^st^ epochs compared to 2^nd^ epochs (Fig. 7e). Finally, we found a significant positive correlation between the total number of wrong answers given (irrespectively of task difficulty) and the total number of non-integrated time-points during the working memory experiment for the #1 subnetwork (r=0.28, p<0.045, Spearman correlation coefficient, see also Fig. 7f).

**Figure 7.**
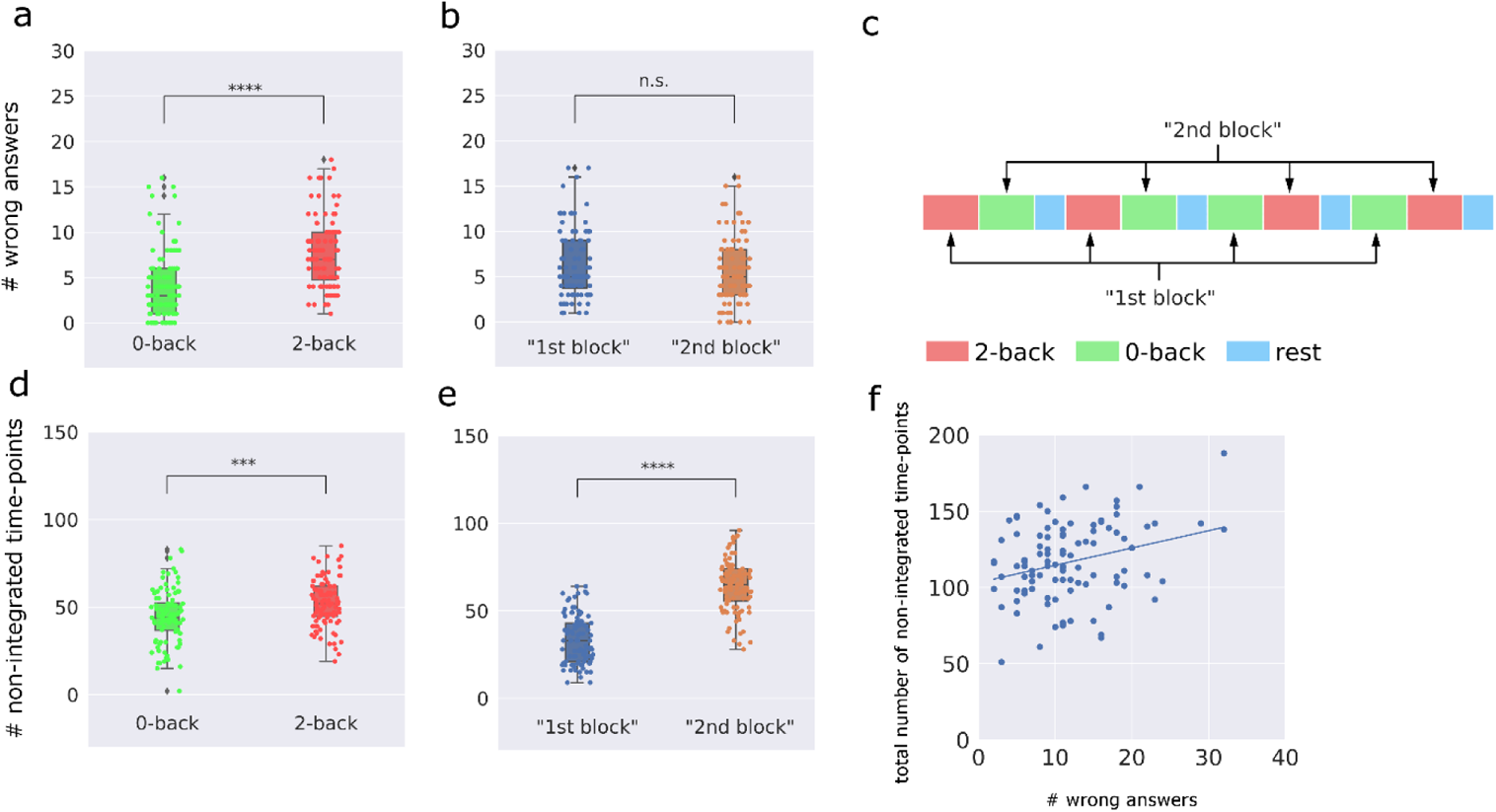
The relationship between degree of integration for the top most integrated subnetwork (SN #1, see also Fig. 3a) and task performance during the working memory experiment. The number of wrong answers given by each subject was significantly larger during the 2-back epochs compared to the 0-back epochs (p<0.0001) (**a**), whereas there was no significant difference between number of wrong responses given during “1^st^ epochs” compared to “2^nd^ epochs” (p=0.3, panel **b**). A pictorial description of “1^st^ epochs” and “2^nd^ epochs” is given in panel (**c**). The number of non-integrated time-points during 2-back epochs was significantly larger than for 0-back epochs (p<0.0001, panel **d**) as well as for 2^nd^ epochs versus 1^st^ epochs (p<0.0001, panel **e**). All statistical tests were performed using permutation tests, 10000 permutations). The graph presented in panel (**f**) shows the correlation between the total number of non-integrated time-points for SN #1 during the entire experiment and the total number of wrong answers given by the participants (r=0.28, p<0.045, Spearman correlation coefficient).

### Subnetwork integration, flexibility and quasicyclicity at lower and higher frequencies

All results presented so far have been based on BOLD signal changes that exist in the 0 – 0.1 Hz frequency range (third intrinsic mode function, IMF 3, peak of power at approx. 0.04 Hz, see also Suppl. Fig. 1b). However, it is clear from Suppl. Figs. 1b-d that in all three experiments, there exists a slower (IMF 4, peak at approx. 0.02 Hz) as well as a faster (IMF 2, peak at approx. 0.1 Hz) BOLD signal component that might be of relevance for our study of spatiotemporally flexible subnetworks. We therefore performed identical analyses (for brevity, only the results from the working memory experiment are shown) and assembled SNCs and SNs for the slower and faster components respectively (see Suppl. Figs. S8-S10). For the faster component, the total number of assembled subnetworks was reduced by approximately 15 percent, whereas the analysis for the slower component yielded a slight increase (approx. 2.5 percent) of subnetworks (Suppl. Figs. S8g and S8h). Importantly, similar to the main results, the majority of the SNs contained very few SNCs and a relatively small number of SNs contained up to several thousand SNCs. For the slower BOLD signal component, the SNs stayed integrated for longer periods of time with fewer instances of integration. For the faster signal component, we observed the opposite behavior with shorter integration periods and a higher frequency of instances of integration (see also Suppl. Figs. S8d and S8i). Although the spatial topology of the five most integrated subnetworks for the slower and faster signal components are not identical to the main results, the majority of included parcels are shared across frequencies, including the visual cortex, dorsal attention network and basal ganglia (Suppl. Fig. S9). Moreover, the degree of spatial diversity and temporal participation at the level of individual parcels share many similarities across frequency bands (Supp. Fig. S10). Together with the findings pertaining to the quasicyclicity of correlation and anti-correlation of subnetwork amplitudes (Suppl. Figs. S8e and S8j), our results suggest a spatial coupling between spatiotemporally flexible subnetworks that resides in different BOLD signal frequency ranges.

The results from our replicability study using the “LR” datasets are shown in Suppl. Figs. S11-S15.

### Global synchronicity and spatial overlap between subnetworks

Hitherto, we have shown that BOLD brain activity signals can be represented and studied as spatiotemporally flexible subnetworks, which are built within a combinatorial framework that includes instantaneous phase synchronicity and modularity as key pillar stones. Further, by focusing on the top five most integrated subnetworks (highest q_int_ values) we have shown that their internal spatial topology is closely related to the activation maps typically found for voxel-based, general linear model studies of brain activation. In the case of rest, spatiotemporally flexible subnetworks show a high degree of spatial similarity with the canonical resting-state networks. Furthermore, we have presented results suggesting that subnetworks, to a varying degree, switch between being internally integrated to being disintegrated and then switch back again. A key question is the following - if we consider all subnetworks and not just the top most integrated ones, how similar is the degree of synchronicity and spatial overlap between them and is this dependent on task? We attempt to answer this question by computing the spatial dice coefficient between all pairs of subnetworks and by computing the pairwise correlation coefficient between phase signal time-courses for all subnetworks (Fig. 8). The latter is a measure of the tendency towards either integration or segregation between distinct subnetworks.

**Figure 8.**
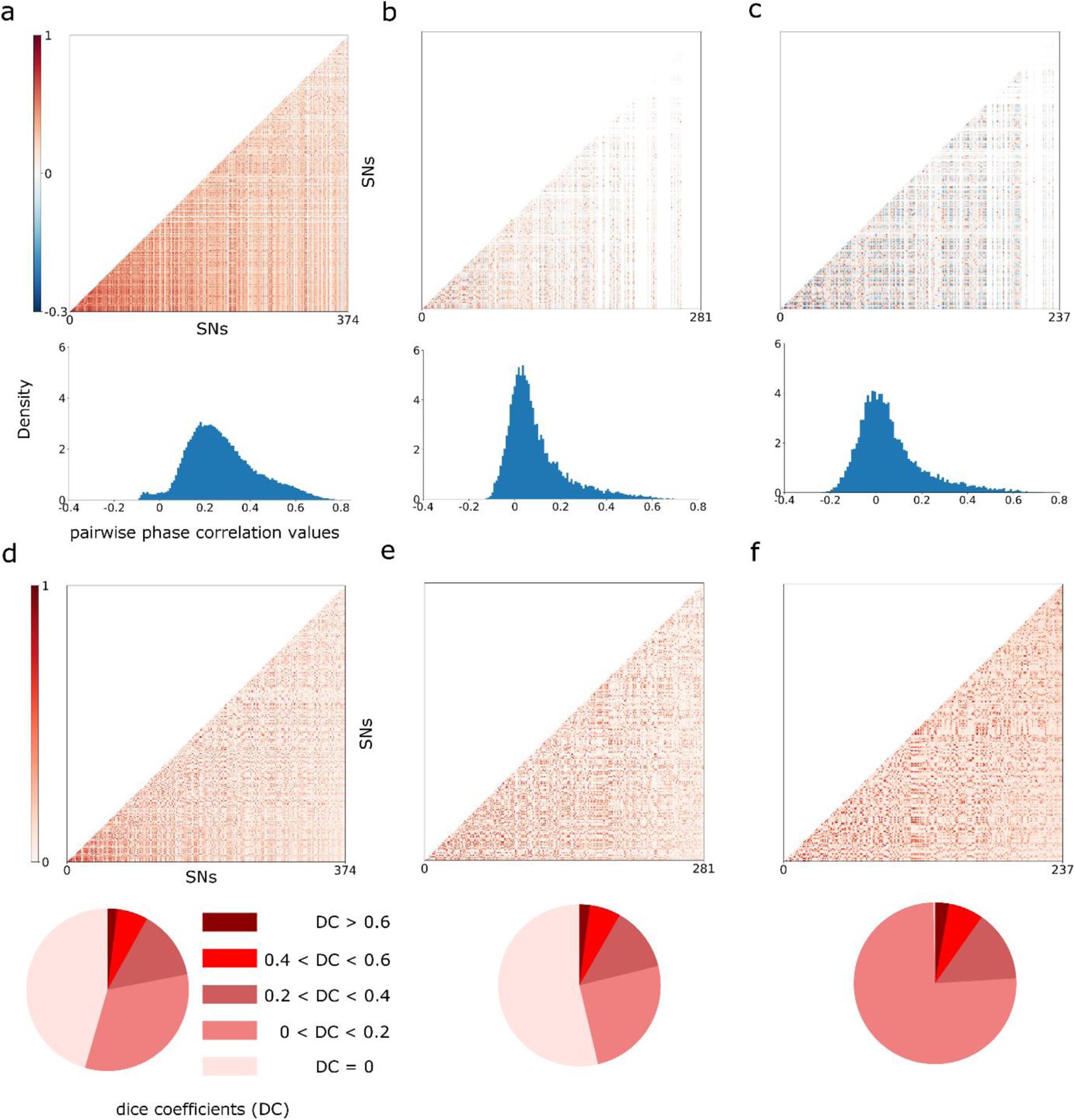
The degree of temporal synchronicity between all pairs of subnetworks shown in the form of correlation matrices and histograms of correlation values for working memory (**a**), motor (**b**) and rest (**c**). Mean correlation values: r = 0.25 (working memory), r = 0.03 (motor), r = 0.04 (rest). The probability density is plotted on the y-axis for the histograms. The lower panel shows the degree of pair-wise spatial overlap (dice coefficient) between all subnetwork for working memory (**d**), motor (**e**) and rest (**f**). Mean of dice coefficients: 0.11 (working memory), 0.10 (motor) and 0.15 (rest). For increased visibility, the relative proportion of dice coefficients (DC) were divided up into five intervals that are shown as pie charts.

The results pertaining to the degree of temporal synchronicity among subnetworks are shown in Figs 8a-c. Starting with the results for rest (Fig. 8c), we note that the peak of the distribution of correlation coefficients between pairwise comparisons of phase signal time-courses is located at 0 with a long positive tail. Similar results were obtained for the motor task (Fig. 8b), albeit with a smaller positive tail and less negative relationships between subnetwork phase time-courses. However, for the memory task, the large majority of subnetwork phase time-courses are positively correlated with each other (Fig. 8a). Indeed, the distribution of subnetwork phase correlations for the working memory experiment was significantly different from both motor (p<0.0001) and rest (p<0.0001) whereas no significant difference was found between motor and rest (p=0.14) (permutation tests, 10000 permutations). This result implies that subnetwork integration was more pronounced for the working memory task compared to the motor and rest experiments.

The degree of spatial overlap among subnetworks assembled during tasks and rest are shown in Fig. 8d-f. The dice coefficient matrices and the corresponding pie charts shows that for the large majority subnetworks, their degree of overlap is quite small (dice coefficient values below 0.2). Moreover, the results show that the relative proportion of highly overlapping versus spatially more segregated subnetworks remain largely intact across tasks, with a notable exception of a slight increase in overlap during rest for the low end tail of spatial overlap (Fig. 8f). The corresponding results for the faster (IMF4) and slower (IMF2) frequency components during the working memory experiments are shown in Suppl. Fig. 16. Taken together, these results suggest that the working memory task impose an overall substantial increase in temporal synchronicity, i.e. integration, for all assembled subnetworks whereas the degree of spatial segregation between individual subnetworks remain relatively unperturbed by task performance compared to rest.

## Discussion

### Task performance entails phase synchronization across subnetworks

The present findings support our hypothesis that integration and segregation among subnetworks are dynamic events that are simultaneously occurring in the brain during the time span of a single fMRI experiment. In particular, our results provide empirical support that individual subnetworks oscillate between periods of integration and disintegration while separate subnetworks oscillate between integration and segregation vis-à-vis each other. In the case of resting-state fMRI recordings, oscillations of integration and disintegration occur spontaneously and unsynchronized with respect to each other, both within subjects and when averaged across subjects (Figs. 3c and 8a). The most integrated subnetworks (i.e. subnetworks with the highest q_int_ values) show a high degree of spatial overlap to previously described canonical resting-state networks derived from static (i.e. averaged across time) measures of brain connectivity (Damoiseaux et al., 2006, Smith et al., 2009).

On the other hand, if epochs of task are incorporated into the fMRI experiment, the formation of subnetworks and their variability in integration across time is consequently altered (Figs. 3a and 3b). During task-based fMRI, individual subnetworks show a strong tendency to be synchronized across time. That is, the subnetwork phase time-courses become dependent on the specific timing of task epochs and the intermittent epochs of rest. The change towards synchronization, i.e. integration, between subnetworks was particularly strong during the working memory task, with marked positive peaks in phase amplitude during the intermittent rest epochs. Importantly, synchronicity in phase is not limited to the top five most integrated subnetworks shown in Fig 3, but permeates to most assembled subnetworks, which is apparent from the observation that the majority of subnetwork phase-time courses are positively correlated with each other (Fig. 8a). These results are in contrast to rest, for which the peak of the distribution of phase correlation values is close to zero (Fig. 8c).

### Integration of subnetworks during rest and tasks

The inclusion of a task during the fMRI experiment introduces a major shift, in particular for the working memory task, towards an overall synchronization in phase between most subnetworks. However, the average duration of time periods of subnetwork integration does not markedly differ between experiments, notwithstanding the notable exception of the top most integrated subnetwork (SN #1) for the working memory experiment that exhibits an almost 3-fold increase in duration of integration periods compared to the average integration time (Fig. 4). The results shown in Suppl. Fig. S6a suggests that the timing of the working memory paradigm exerts a strong influence on when subnetworks tend to be integrated versus being disintegrated. Overall, subnetworks are most integrated (across subjects) for time periods that starts during the intermittent rest epochs and approximately end after the termination of the succeeding 0/2-back task epoch.

Our results may at a first glance considered to be contrary to the results presented in our previous study that showed strong peaks of network segregation and reconfiguration during the intermittent epochs of rest (Fransson et al., 2018). However, our previous findings and the current results do not contradict each other if we carefully consider the differences in methodological design and the implications thereof. First, in Fransson et al., 2018 we used a temporally static division of brain parcels into networks (following the parcellation scheme provided in Power et al., 2011). Second, we used a point-wise estimates of network integration/segregation (Segregation and Integration Difference, SID) that first sums up within-canonical network node strength and subsequently subtracts between-canonical network node strength. The previously reported segregation during rest epochs should be interpreted vis-à-vis statically defined subnetworks. Moreover, an increase in SID network segregation can be driven by either an increase in within-canonical network connectivity or a decrease in between-canonical network connectivity or a combination of both. Thus, taking the present results into account, the previously documented increase in SID segregation is likely driven by an increased within-network connectivity for the canonical visual network that is included in the most integrated spatiotemporally flexible subnetworks.

Our current results suggest that the organization of spatiotemporally flexible networks exhibits quasi-cyclic periods of integration and disintegration, a characteristic that is not exclusive to rest as previously shown (Strindberg et al., 2021), but also occur during task performance. In this context, it is interesting to contemplate the scenario that the temporal patterns of subnetwork integration during task performance might be of equal importance to understand brain function compared to measuring relative increases/decreases in BOLD signal activity.

### The temporal structure of task-based fMRI experiments and its relationship to spatiotemporally flexible subnetworks

The finding that subnetworks tend to be most integrated during the intermittent rest epochs and the trailing 0/2-back epoch in the working memory experiment is notable (Suppl. Fig. S6). In this context, it is noteworthy that the recurrence plots (Fig. 6a) display a strong pattern of global quasicyclicity during the same time intervals, i.e. a back-and-forth switch between correlation and anticorrelation in amplitude between subnetworks. This observation is paralleled by a marked increase in size (i.e. number of participating parcels) for the most integrated subnetworks (Fig. 6d). Additionally, as described earlier, we observed a strong positive peak in subnetwork phase during the intermittent rest epochs. Interestingly, the motor fMRI experiment did not show similarly marked differences in subnetwork integration, global cyclicity and increases in subnetwork size tied to the temporal structure of the experiment. Why the presence of intermittent rest epochs seems to have a profound importance for the spatial as well as temporal evolution of subnetworks during the working memory experiment is presently unclear. However, in this context it is worth pointing out that current theories for working memory processes point towards the importance of interactions between different neural oscillatory mechanisms in different cortical layers (Miller et al., 2020). Future studies using the methodology presented here applied to MEG data acquired during working memory tasks could potential be able to provide further insights. Nevertheless, our findings give credence to the idea that neuronal processes occurring during intermittent baseline or rest epochs, which are commonly employed in task-based fMRI experiments, play an essential part for succeeding tasks to be carried our according to prior instructions and adapting to changing cognitive demands. In this context, it is important to remember that our method ranks and analyze changes in network properties at a global time-scale (i.e. for the entirety of the fMRI experiment by computing the parameter q_int_).

### Subnetwork synchronization and whole-brain level activation

It is enlightening to relate our findings to those described in an earlier study that used dense sampling (data from 100 fMRI runs collected in three subjects) to increase the signal to-noise ratio by a factor of six in a task-based fMRI paradigm that included a visual checker-board stimulation combined with a letter/number discrimination task (Gonzalez-Castillo et al., 2012). Remarkably, the authors could show that up to 96 % of all voxels in the brain were in a systematical way activated for the task, although showing variability in terms of shapes of BOLD responses. The idea that task-induced modulations of BOLD responses are present at a whole-brain level resonates well with our network-based results. That is, we hypothesize that the increase in change in activity for almost all voxels in response to a task as described in Gonzalez-Castillo et al., 2012 is a reflection of the general increases in synchronization (with respect to the temporal timing of the task paradigm) among subnetworks, as well as increases in amplitude quasicyclicity and periods of increased integration during task performance.

### Spatiotemporally flexible subnetworks and states of brain activity

A contemporary aspect of functional brain connectivity research is the concept of “states”, which has come to play a central role when analyzing network properties that change over time (Allen et al., 2014; Vidaurre et al., 2017; Shine et al., 2017; van der Meer et al., 2020). Different methods have been proposed to estimate brain states during resting-state, ranging from Hidden Markov Models (HMMs) (Vidaurre et al., 2017), k-mean clustering of sliding window co-variance data (Allen et al., 2014), k-mean clustering of time-frequency datasets (Yaesoubi et al., 2015) to “cartographic profiling” (Shine et al., 2016). Although the mentioned approaches make use of different assumptions and claims of whether transitions between states are “smooth” or occur instantaneously, i.e. from one sample point to another, it is difficult to enforce the postulation of distinct brain states onto our results. Although a temporal clustering of the results presented here may in principle be used to define brain states, we do not think that such an approach would provide any further mechanistic insight. In fact, due to the global perspective in the temporal domain that is embedded into our methodological approach (i.e. q_int_), a clustering based on, e.g. subnetwork phase (i.e. degree of integration), spatial flexibility or combinations thereof, would therefore produce states that are unlikely to reflect distinct and momentary changes in neuronal processing.

Rather than reasoning in terms of states, our approach is in alignment with the concept of quasiperiodic patterns of oscillatory brain activity as described previously (Majeed et al., 2011). In recent study, the same group demonstrated that quasiperiodic patterns propagate along macroscopic gradients in the brain (Yousefi and Keilholtz, 2021) Interestingly, in another recent study that used the same input data as used here (Abbas et al., 2019) but focused on quasi-periodical oscillations between the task positive and default mode networks (Fransson 2005, Fox et al., 2005), the authors could show that the overall pattern of spontaneous fluctuations was significantly altered during task compared to rest. Although the Abbas study largely treated quasiperiodic signal changes as an additive signal component that optionally can be removed from the data if needed, the results in the Abbas study is in agreement with results presented here with regard to the fact that task and/or spontaneously driven oscillatory activity is present during task performance. Of note, the presence of spontaneous activity during the performance of a 2-back working memory task, that employed a sustained paradigm design, was early on shown to exist in the default mode and task positive networks (Fransson, 2006).

Additionally, further insights may be gained from studying the temporal evolution of the phase amplitudes for individual parcels in the most integrated subnetworks as exemplified in Supplementary Movies M1-M3. These movies suggest that the quasicyclical nature of subnetwork activity starts with activity in a single parcel that is followed by a gradual recruitment of additional parcels until a peak is reached, which is then in turn succeeded by a gradual decrease of participating parcels. This repetitive pattern is present throughout the entire fMRI experiment, which suggests that a quasicyclic behavior which implies a repetitive process of gradual build-up and dismantlement of subnetworks is a core feature of brain activity.

### Temporal ordering of data in time-resolved analysis of network brain connectivity

Commonly used methods to derive network-based models of time-resolved functional connectivity such as sliding window and time-frequency based methods do retain the temporal order of the data. Here we used the instantaneous phase synchronization analysis which also adhere to the principle of keeping the temporal order of the data. However, alternatives strategies to compute functional connectivity has been proposed and used extensively in the literature. For example, using the concept of co-activation patterns (CAPs, Liu and Duyn, 2011, Liu et al., 2018; Karahanoglu and Van De Ville, 2018) brain connectivity at individual time-points is modelled, but the temporal order is subsequently discarded when clustering over non-continuous data points. Similarly, recent studies focusing on so called edge-centric connectivity analysis combined with clustering methods (Faskowitz et al., 2020) have suggested that events of high-amplitude co-fluctuations synchronizes across subjects during movie watching and that they carry substantially more information compared to low-amplitude events (Esfahlani et al., 2020). There are clearly several benefits from using such methods, in particular in terms of data dimensionality reduction to extract properties that are consistent and interesting from an inter-subject variability versus personal traits perspective. However, based on the results presented here, we think that preserving the temporal ordering of the data-points in the analysis provides additional benefits if a fuller picture of the temporal evolution of network changes across time is desired.

### Multi-scale behavior of spatiotemporal networks

It is well known that the spatial topology of spontaneous resting-state network activity and the corresponding static functional networks are dependent on the frequency range of BOLD signals (Niazy et al., 2011; Thompson & Fransson, 2015). Although the potential of addressing the temporal multi-scale properties of network behavior has recently been discussed (Betzel and Bassett, 2017), it has to our knowledge not previously been directly addressed in a time-resolved fMRI brain connectivity setting. Given the prerequisites by the EMD algorithm, our results show that spatiotemporally flexible subnetworks can be assembled and studied in different frequency bands (Suppl. Figs. S1, S8-9 and S16). Interestingly, subnetworks that are strongly integrated across time shows a large degree of spatial overlap across time-scales (compare e.g. Fig. 3 with Suppl. Fig S9). This finding implies that although time-resolved networks show periodic changes in integration versus disintegration that are apparent in different frequency bands, they mutually show a substantial degree of spatial overlap. Obviously, this observation does not exclude the possibility of other methods beside the EMD algorithm to assess frequency-specific BOLD signal changes might reveal significant cross-frequency differences in spatial overlap between networks.

It is of interest to compare properties of oscillating, quasiperiodic subnetworks based on hemodynamic signal changes reported here to recent results obtained from intra-cortical electrical measurements (Mostame et al., 2019, 2020; Sadaghiani et al., 2022). Although methods to estimate connectivity are different and the spatial coverage is more restricted, ECoG measurement of both tasks and rest suggests that fast intrinsic oscillations are dominated by a spatial organization that are shared across frequency bands (Mostame et al., 2021). This finding is in agreement with our findings in the BOLD regime, in so far that the spatial topology of highly integrated subnetworks shows similarities across frequency bands (Suppl. Fig. S9). Relatedly, the authors reported that cognitive state (e.g. a working memory task) did not in any significant manner influence oscillatory activity, a finding which may appear to contradict the state-dependent re-organization of spatiotemporally flexible subnetworks reported here. However, these discrepant observations may be attributed to differences in analysis methods (i.e. phase coupling versus correlation in instantaneous phase changes). Whereas Mostame et al. focused their analysis on time-locked averaging of data recorded during brief pre and post-stimuli periods, our analysis was designed to be sensitive to signal changes for the fMRI experiment in its entirety. While we do not claim to be able to unequivocally distinguish between simultaneously ongoing intrinsic (i.e. task-independent) oscillations and task-evoked responses, we suggest that our results show that strong interactions occur between the two during task-based fMRI.

### Limitations

Like most other methods that aim to compute time-resolved measures of brain network connectivity from fMRI data, concerns regarding putative methodological caveats and user-specified choices of parameters should be addressed and discussed. To that end, our method uses the Louvain algorithm, which is known to not provide identical divisions of the data into communities across separate runs. Although we ran the algorithm 100 times to achieve a consensus with regard to community membership, it cannot be ruled out that other algorithms would produce improved results (e.g. the Infomap algorithm described in Rosvall et al., 2009).

Moreover, the iterative process of assembling SNC from the seed pairs is dependent on the spatial granularity of the parcellation scheme used (see also recent paper on the effect of choice of parcellation scheme, Salehi et al., 2020). A more fine-grained parcellation would require SNC to be larger than used here. Additionally, we used an expansion factor of ten, meaning that for each seed pair, the top ten pairs (i.e. pairs with the highest q_int_ values) were kept at each iteration. The value of the expansion factor was set arbitrarily, but we have in our previous study studied the effect of increasing the value of the expansion factor, see Supplementary Material in Strindberg et al., 2021 for additional results and discussion. Further, the decision of when SNCs are considered to be collectively integrated and thus merged into a SNs employed the use of a heuristically defined parameter OV (the co called “no-split” criteria, supplementary methods for further details).

Notably, we used the empirical mode decomposition algorithm to separate signal variability into different frequency bands. Although this method has previously been applied in the context of BOLD fMRI data (e.g. Niazy et al., 2011), the most favored method in the fMRI literature to accomplish this goal is to use bandpass filters. Quantitative changes between the filtering approaches were addressed in our previous study that included a direct comparison between the empirical mode decomposition method and bandpass filtering of resting-state BOLD signals, which showed a high correspondence between them (see Suppl. Figs. S29-S34 in Strindberg et al., 2021).

### Conclusions and future directions

Using a bottom-up approach to represent time-resolved functional network connectivity, starting with pairs of parcels that in a step-wise manner are assembled into subnetwork components and finally merged into subnetworks, we have demonstrated quasi-periodic properties of subnetwork integration, disintegration and segregation at a whole-brain level. Our results highlight the importance of oscillations in fMRI network connectivity that transcends beyond resting-state conditions to include task-based paradigms that are typically employed in functional neuroimaging research. Although further investigations employing other tasks and different temporal experimental designs needs be carried out, the results presented here suggest that a hierarchical and data-driven analytical approach delivering quantitative information on network integration, flexibility and quasicyclicity provides valuable insights into brain function that is not otherwise readily accessible. Indeed, the importance of oscillations in network activity during tasks has increasingly been recognized as an important factor for modelling spiking neuronal activity related to memory replay and multisensory cues (Korcsak-Gorzo et al., 2022). Thus, an expose of network rhythmicity that takes into account the fMRI experiment in its entirety as shown in the current work is of relevance to fully understand the relationship between cognitive work and brain network activity.

## Acknowledgements

P.F. was supported by the Swedish Research Council (grant No. 2020-03288) and the Swedish e-Science Research Center. M.S. was supported by Jerring Foundation and Sällskapet Barnavård. Data were provided by the Human Connectome Project, WU-Minn Consortium (Principal Investigators: David Van Essen and Kamil Ugurbil; 1U54MH091657) funded by the 16 NIH Institutes and Centers that support the NIH Blueprint for Neuroscience Research; and by the McDonnell Center for Systems Neuroscience at Washington University. The funders had no role in study design, data collection and analysis, decision to publish, or preparation of the manuscript.

## Code availability

All code used in the present work is available at: https://github.com/MarikaStrindberg/TimeResolvedNetworks DOI:10.5281/zendo.4794699

## Author contributions

Conceptualization: PF, MS; Methodology: PF, MS; Software: PF, MS; Visualization: PF, MS; Writing - original draft: PF, MS; Writing – Review & Editing: PF, MS

