## Supplementary material for "Brain network integration, flexibility and quasicyclicity during task and rest": Suppl. Methods and Figures

Supplementary methods and figures for Fransson & Strindberg, 2022.

19-APRIL-2022

*Supplementary Methods*

*Data used.*

The 100 independent subject dataset from the Human Connectome 500 subject release was used (van Essen et al., 2012; Smith et al., 2013). For our primary, discovery analysis, "RL", i.e. phase-encoding gradient in the right-left direction. A replicability study of our results was performed on the "LR". Further information regarding the MR acquisition parameters can be found in (Uğurbil et al., 2013). The fMRI data had undergone image preprocessing and removal of artifacts from non-neuronal origins by means of the FIX ICA (FMRIB's Independent Component Analysis-based X-noisifier) data artifact rejection process which removed ICA components from the data that were considered to constitute signal contributions from white matter, cerebro-spinal fluid, head movement, cardiac and respiratory sources (Gasser et al., 2013; Griffanti et al., 2014; Salimi-Khorshidi). The data consisted of 2x1200 (REST) and 2x405 (WM) image volumes per subject (TR = 0.72). To facilitate a fair comparison between rest and tasks, and in particular the working memory task, only the first 405 time-points of the resting-state recordings were used in the analysis.

*Experimental paradigms.*

The working memory task employed a version of the n-back working memory task where 2-back and 0-back epochs (10 trials in each epoch, duration 25 seconds) were interleaved with epoch of baseline condition (15 seconds). Each session contained 8 task epochs (4 of each type) where each epoch contained trials that consisted of visual presentations of objects of "faces", "body parts", "tools" and "places". Further information regarding the details for the stimuli presentation and paradigm design is found in Barch et al., 2013.

*Assembling subnetwork components (SNCs).*

The algorithm to find gradually larger SNCs started by identifying seed pairs as described in the main text. The seed pairs represented the pair of highest pairwise $q_{int}$ from the perspective of each area in the parcellation. At each iteration, the relative frequency ($q_{int}$) to which each pair was assigned to the same community was computed for all possible pairwise combinations of parcels. Due to the fact that only the most frequent constellations from the point of view of individual parcels were of interest, an expansion factor had to be set for each iteration in the algorithm. An expansion factor K = 10 parcels was used here. This setting implies that for each seed pair, the top (i.e. highest $q_{int}$ values) ten pairs of parcels of which the seed pair was most frequently found to be assigned to the same community with were kept. The ten resulting SNCs of size n = 4 (brain parcels) for each seed pair where then used as seeds in the next iterative step where the same process was repeated to find SNCs of size n = 6 etc. For a discussion on effect of variations in the K-parameter and comparisons to randomly generated SNCs, please see Supplementary Material in Strindberg et al., 2021.

*Merging subnetwork components (SNCs) into subnetworks (SNs).*

As previously described (Strindberg et al., 2021), we adhere to two principles when merging SNCs into SNs. First, the most integrated (averaged across subjects) SNC across time was ranked as number one and named SN #1. Second, all other SNCs that shared at least 5 parcels (out of the maximum of 8 areas) was merged together in SN #1. Subsequently, all SNCs incorporated into SN #1 was removed from the list of SNCs and the process to merge SNCs was iterated until all SNCs had been assigned to a SN (for an illustrative presentation, see Suppl. Fig. S7 in Strindberg et al., 2021). The spatial requirement of 5 shared parcels was heuristically determined in Strindberg et al., 2021. It was based on the criteria that the SNCs representing a particular SN at each time point should be assigned to the same community, i.e. the SN should be internally coherent at all time-point and subjects. Generally, a lower setting of the number of required shared parcels will result in a higher number of SNs that to a larger degree are spatially overlapping with each other. In contrast, a more relaxed constraint of spatial overlap between SNCs will result in fewer SNs. However, in the latter case the criteria of internal coherence is not always fulfilled.

**Supplementary Movie M1**. (SN #1, working memory, Suppl_Movie_M1_wm_fr5.avi) Shows the mean (across SNCs) phase amplitude for all participating brain parcels at each time point during the working memory experiment in a single subject (same subject for all three movies). The color of a square positioned in the center of the screen marks the presence of rest (blue), 0-back (green) or 2-back (red) epochs. The frame rate was set to 5TR/sec.

**Supplementary Movie M2**. (SN #1, motor task, Suppl_Movie_M2_motor_fr5.avi) Shows the mean (across SNCs) phase amplitude for all participating brain parcels at each time point during the motor task experiment in a single subject. The color of the square in the center of the screen marks the presence of rest (blue), tongue (red) or finger (orange) or toes (cyan) movement epochs. The frame rate was set to 5TR/sec.

**Supplementary Movie M2**. (SN #1, rest, Suppl_Movie_M3_rest_fr5.avi) Shows the mean (across SNCs) phase amplitude for all participating brain parcels at each time point during the resting-state experiment in a single subject. A blue square positioned in the center of the screen marks the presence of rest. The frame rate was set to 5TR/sec.

**Supplementary Figures:**

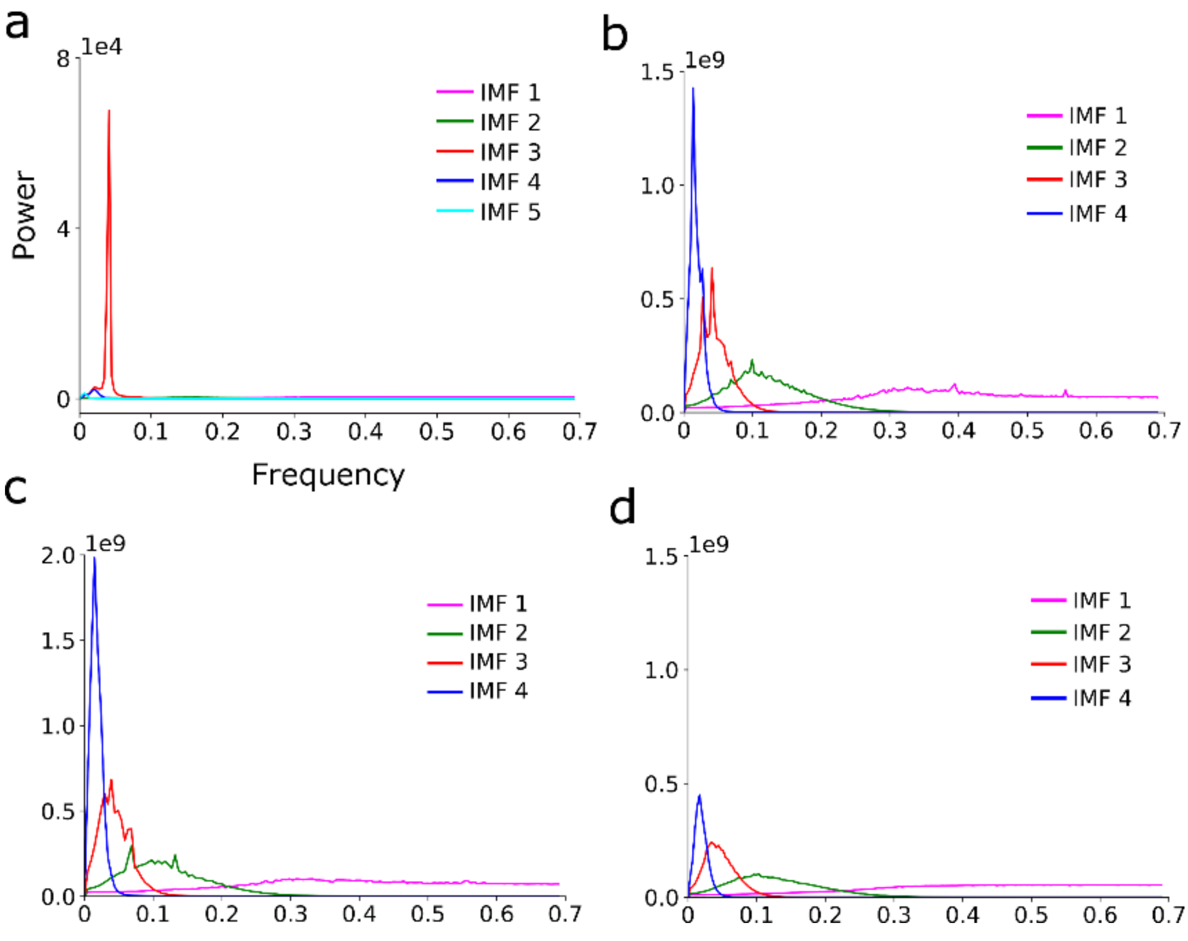


***Supplementary Figure S1.*** *Mean power spectra for intrinsic mode functions (IMFs) averaged across all subjects and parcels for the synthetic data example (**a**) (see also Figure 1 in main text) working memory (**b**), motor (**c**) and rest (**d**). For the main analysis, we*

*selected the third (IMF 3) component in all cases since its frequency range closely matches the low-frequency range of spontaneous as well as block task induced BOLD signal changes (but see also results for IMF2 and IMF4 in Suppl. Figs. S7-9 and S15.*

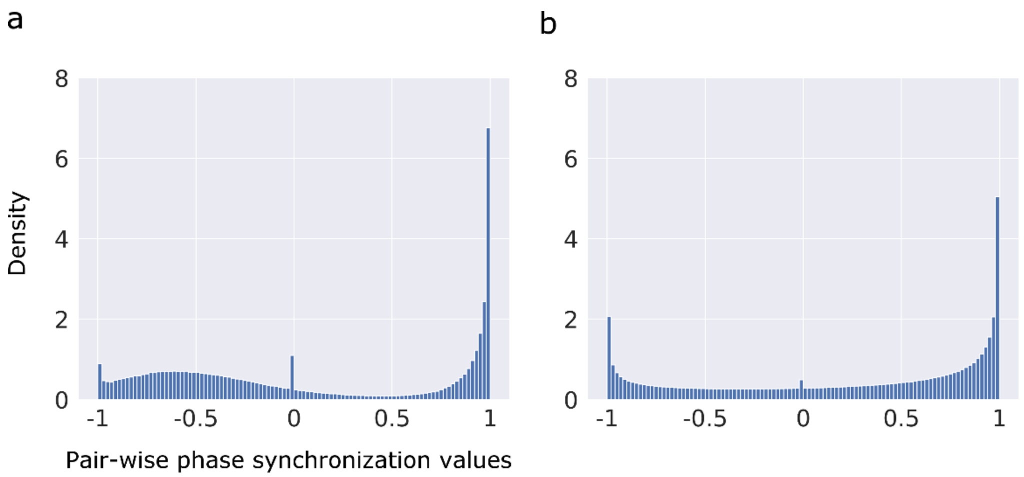


***Supplementary Figure S2**. Histogram of pair-wise phase synchronization values computed for one of the synthetic datasets (**a**) and in one subject across all 264 brain parcels for the working memory task (**b**). Three peaks of phase synchronization values are present at -1, 0 and 1. These peaks were observed in all subjects.*

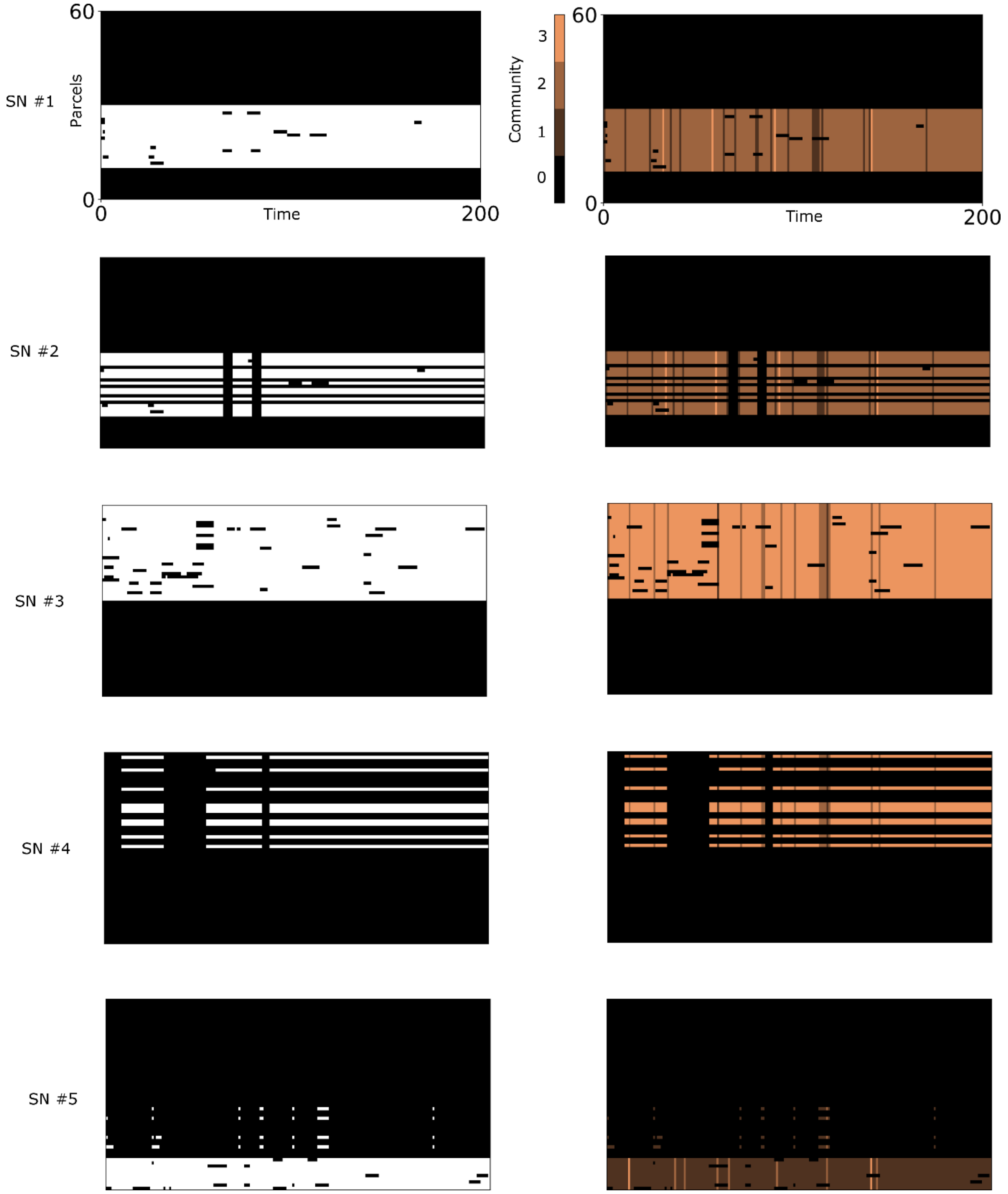


***Supplementary Figure S3****. Example of spatiotemporally flexible properties of subnetworks (SN) in one data set for the toy data example shown in Fig. 1. The left column shows as a function of time which parcels that are included (marked in white) and not included (marked in black) for the five subnetworks. Note that due to the presence of noise, the subnetworks do not perfectly capture the three subsets of parcels shown in Fig. 1a at all points in time. The right column shows the corresponding community membership for each subnetwork as a function of*

*time. Note that a community membership equal to zero implies that the subnetwork is dis-integrated at the given point in time (for parcels that are included in the subnetwork, see left column).*

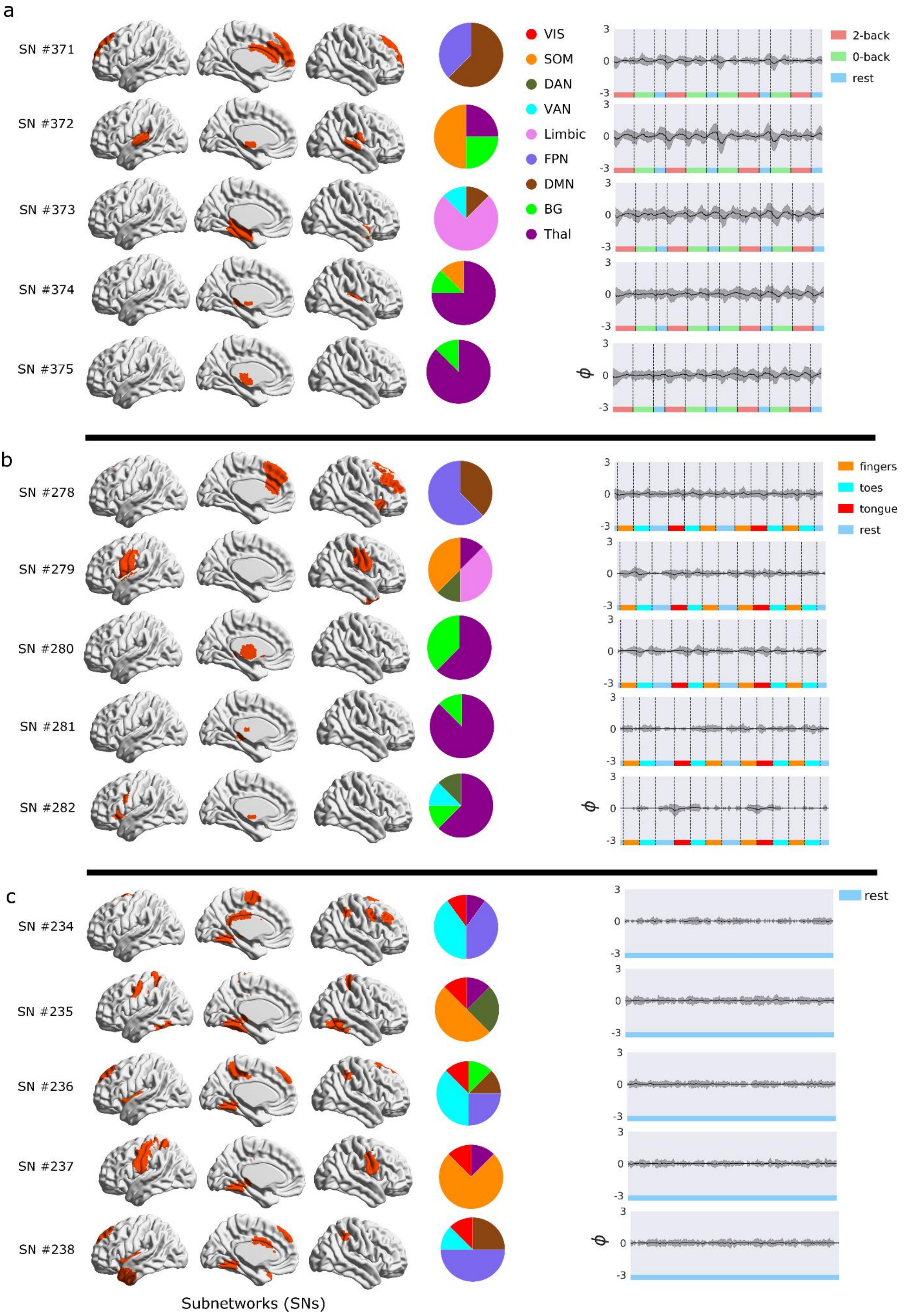

a
SN #371
SN #372
SN #373
SN #374
SN #375
VIS
SOM
DAN
VAN
Limbic
FPN
DMN
BG
Thal
2-back
0-back
rest
3
0
-3
ϕ
b
SN #278
SN #279
SN #280
SN #281
SN #282
fingers
toes
tongue
rest
c
SN #234
SN #235
SN #236
SN #237
SN #238
rest
Subnetworks (SNs)

***Supplementary Figure S4.*** *Spatiotemporally flexible subnetworks (SNs) assembled during working memory (**a**), motor (**b**) and rest (**c**). This figure shows the five least integrated SNs (i.e. the five SNs that had the lowest mean degree of integration ($q_{int}$)). The pie charts in the middle column show the relative contribution (dice coefficient) from the nine canonical resting-state networks to each SN. The graphs to the right show the average (across subjects) phase time-course. For the working memory task (**a**), the least integrated SNs involved thalamus (SN #374, #375), limbic (SN #373), somatomotor (SN #372) and default mode (SN #371) networks (mean $q_{int}$ = 0.016, 0.014, 0.012, 0.012, 0.012; all SNs consisted of a single SNC). During the motor task (**b**), least integrated SNs involved thalamus (SN #280, #281, #282), somatomotor (SN #279) and frontoparietal (SN #278) networks (mean $q_{int}$ = 0.023, 0.020, 0.020, 0.018, 0.016; all SNs consisted of a single SNC). During rest (**c**), the least integrated SNs included parcels located in the somatomotor (SN #236, #237), ventral attention (SN #234, #236) and frontoparietal networks (SN #234, #238) (mean $q_{int}$ = 0.022, 0.021, 0.020, 0.020, 0.019, number of SNCs 3, 1, 1, 1 and 1).*

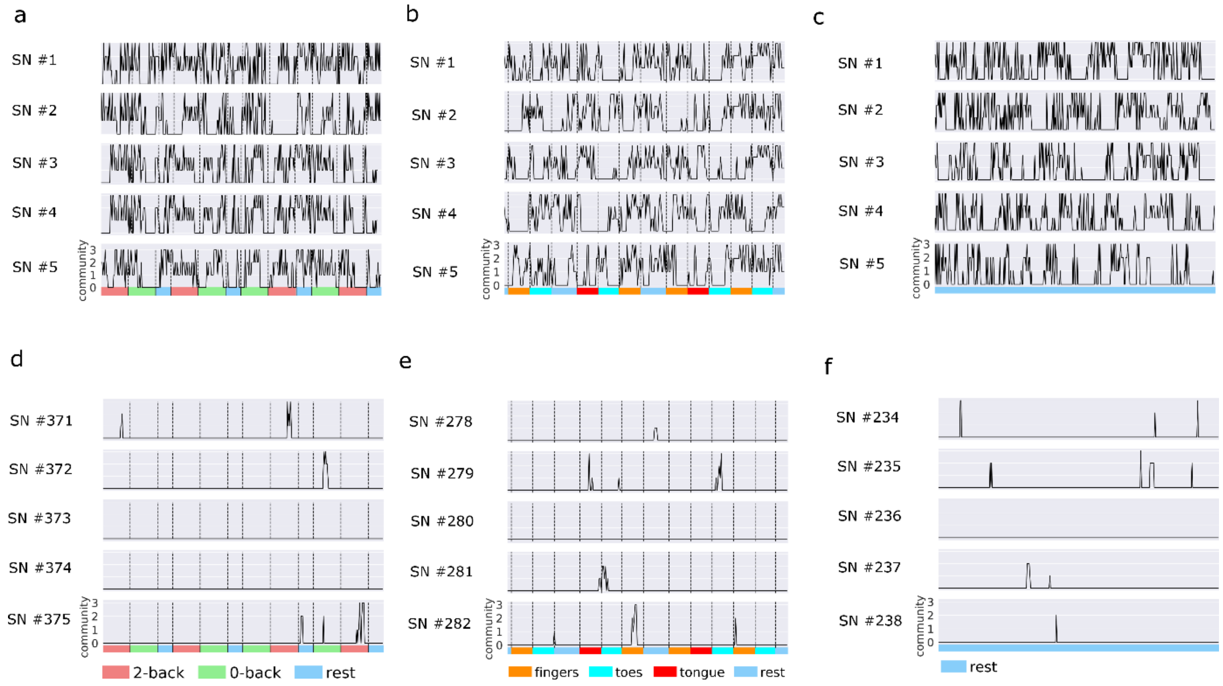


***Supplementary Figure S5.*** *Time-courses for subnetwork (SN) community assignment in a single subject during working memory (**a**,**d**), motor (**b**,**e**) and rest (**c**,**f**). The upper row (**a**,**b**,**c**) shows community assignment for the five most integrated SNs whereas the lower row (**d**,**e**,**f**) display the five least integrated SNs. Note that a community assignment of*

*zero imply that the SN is disintegrated (i.e. none of the SNCs that are incorporated in the SN is integrated at the time).*

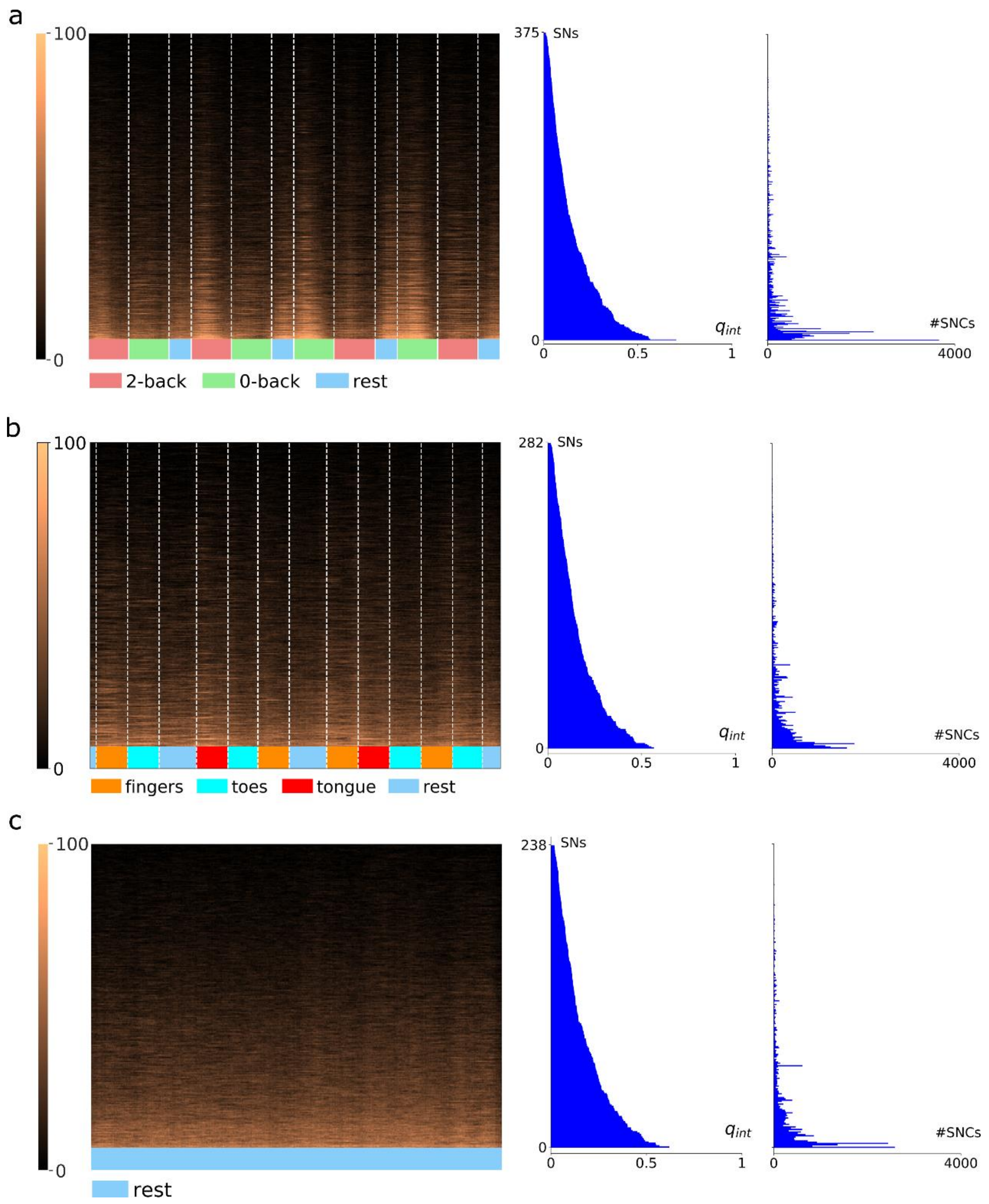


***Supplementary Figure S6****. Degree of integration across subjects (100 = integrated in all subjects) plotted for all subnetworks (SNs) as a function of time during the working memory (**a**), motor (**b**) and rest (**c**) experiments. The graphs in the middle and right columns show the mean $q_{int}$ values (across subjects) and the number of Subnetwork components (SNCs) for each SN, respectively.*

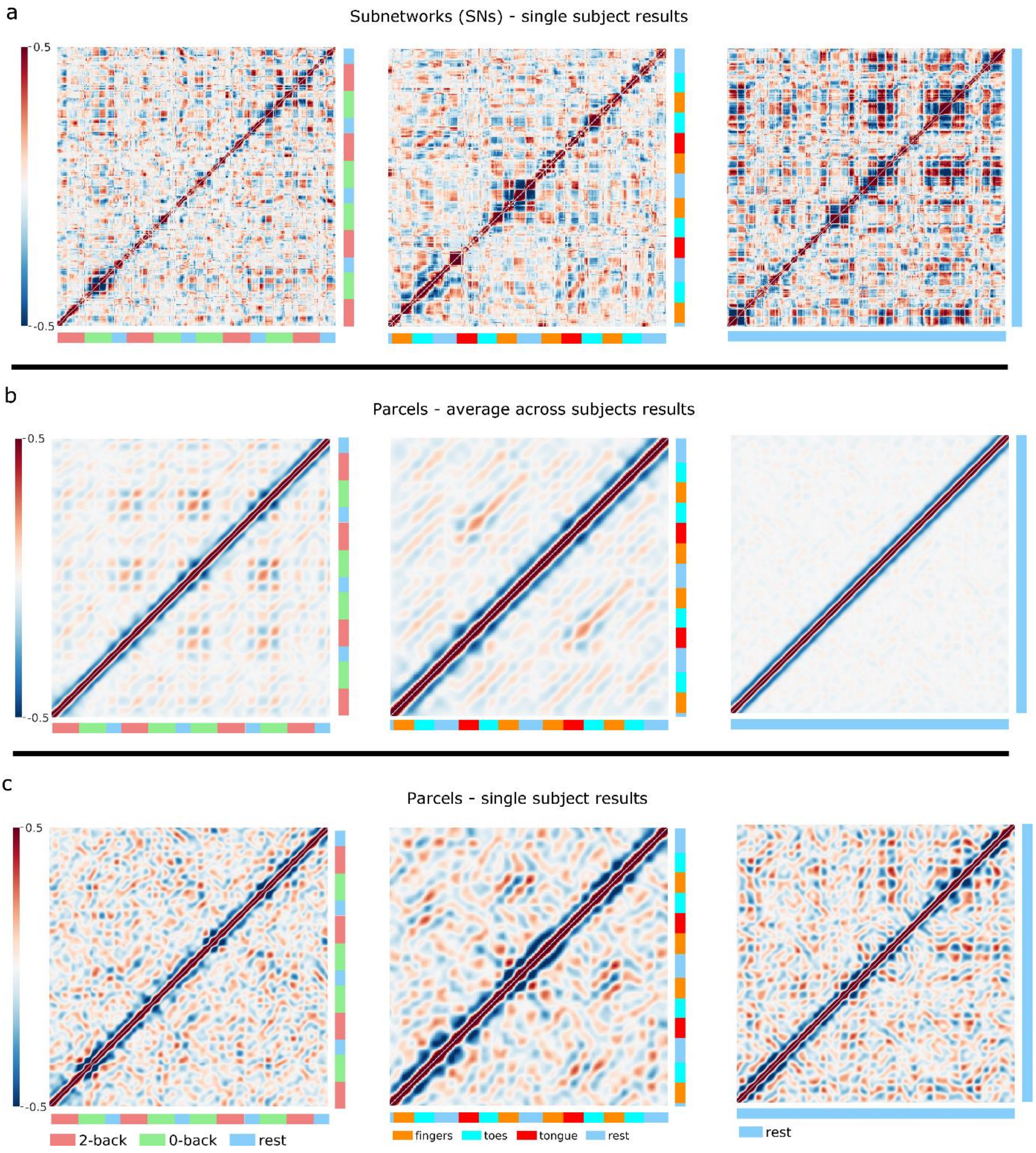


***Supplementary Figure S7****. Recurrence matrices that shows the degree of correlation in amplitude activation and de-activation across subnetworks (SNs) during working memory (left column), motor task (middle column) and rest (right column) at different points in*

*time for a single subject (**a**). Panel (**b**) shows recurrence matrices computed at the level of individual parcels (averaged across subjects) and panel (**c**) shows an example of results at the level of individual parcels in a single subject.*

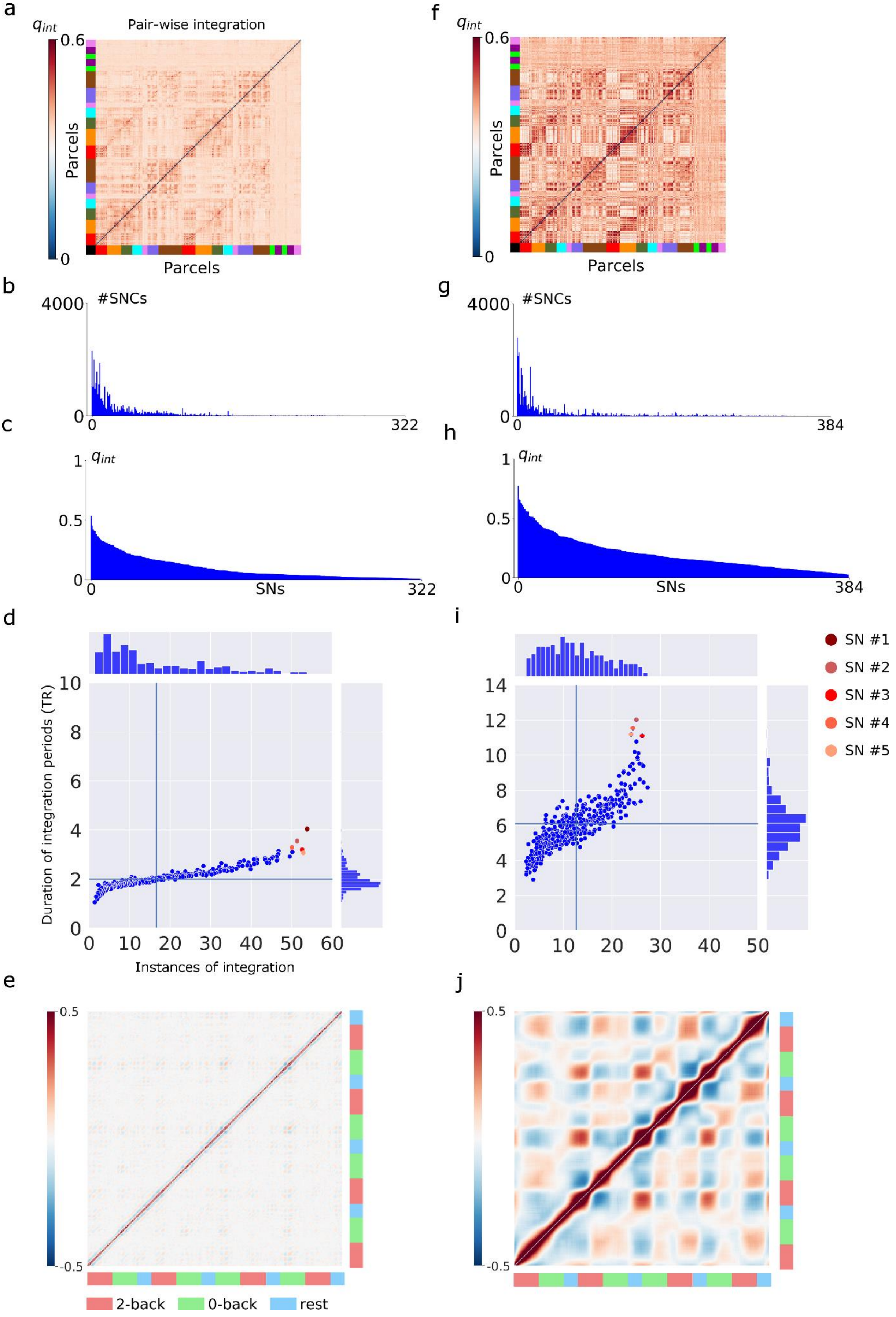
a
f
Pair-wise integration
$q_{int}$
0.6
0
Parcels
Parcels
b
g
#SNCs
4000
0
322
384
c
h
$q_{int}$
1
0.5
SNs
d
i
Duration of integration periods (TR)
Instances of integration
SN #1
SN #2
SN #3
SN #4
SN #5
e
j
0.5
-0.5
2-back
0-back
rest

***Supplementary Figure S8***. *Results for the higher frequency component (IMF2, left column) and lower frequency component (IMF4, right column) (see also Suppl. Fig. S1). For the higher frequency component, the mean duration of integration for SNs was 2.0 TRs and the average number of integration periods was 16.6 (**d**). The corresponding results for the slower BOLD component (IMF4) was 6.1 TRs and on average 12.6 instances of integration during the working memory experiment (**i**). Recurrence matrices of correlation in amplitude activation and deactivation across Subnetworks are shown in panels (**e**) (IMF2) and (**j**) (IMF4).*

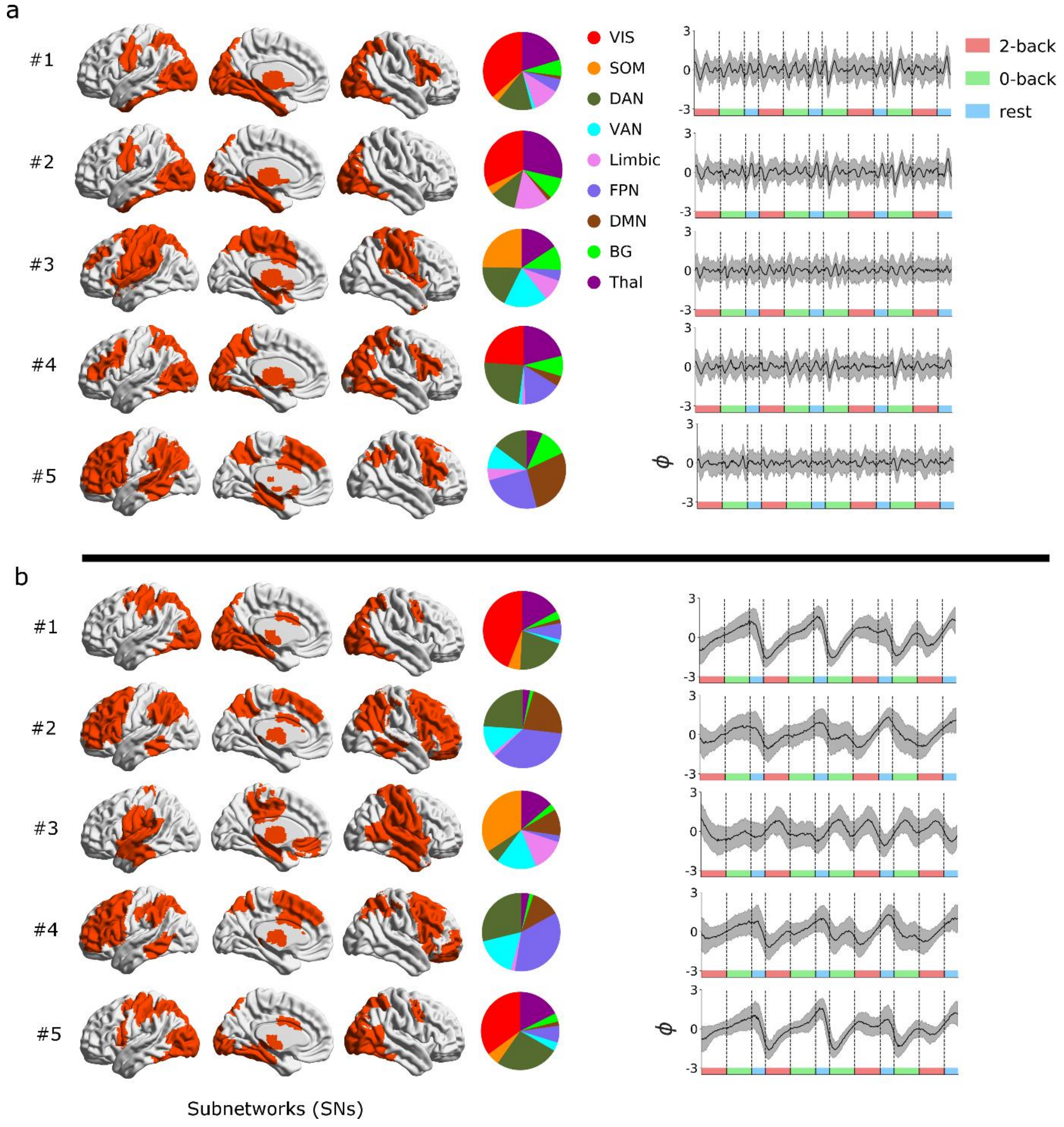

***Supplementary Figure S9****. Spatiotemporally flexible subnetworks (SNs) assembled during the working memory experiment that are computed from the higher frequency BOLD signal component (IMF2, panel* ***a****) and lower frequency signal component (IMF4, panel* ***b****) (see also Suppl. Fig. S1)*

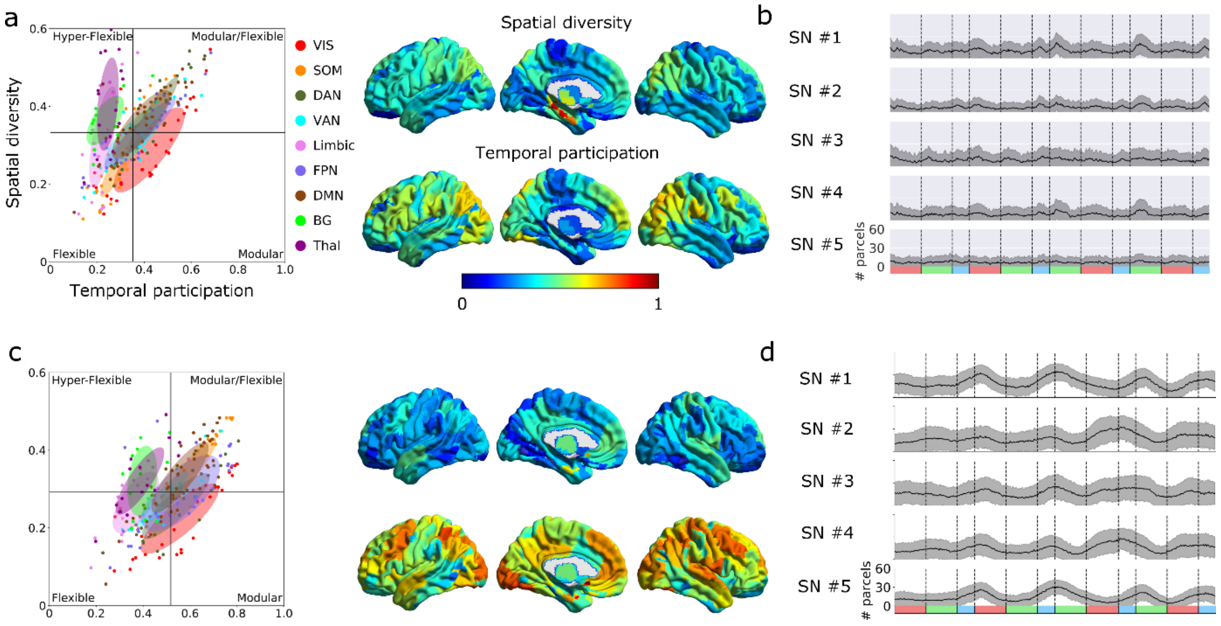


***Supplementary Figure S10****. Spatial diversity and temporal participation calculated at the level of individual parcels for the higher frequency signal component (IMF2, panels* ***a*** *and* ***b****) and lower frequency signal component (IMF4, panels* ***c*** *and* ***d****) (see also Suppl. Fig. S1).*

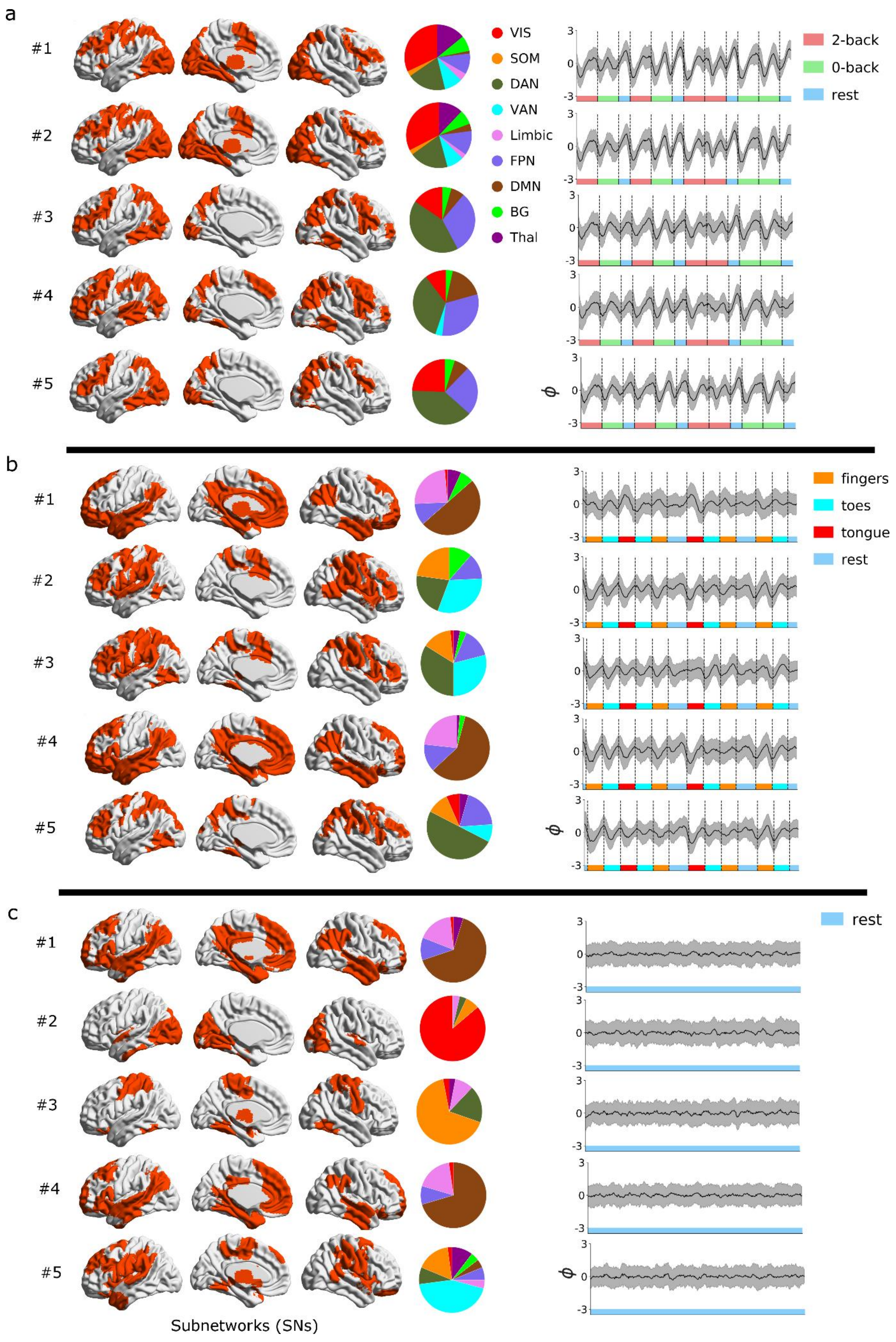
a
#1
#2
#3
#4
#5
VIS
SOM
DAN
VAN
Limbic
FPN
DMN
BG
Thal
2-back
0-back
rest
$\phi$
b
fingers
toes
tongue
rest
c
rest
Subnetworks (SNs)

***Supplementary Figure S11.*** *This figure complements the discovery results shown in Fig. 3 in the main text. The results shown here represents a replication analysis conducted on the "LR" datasets.*

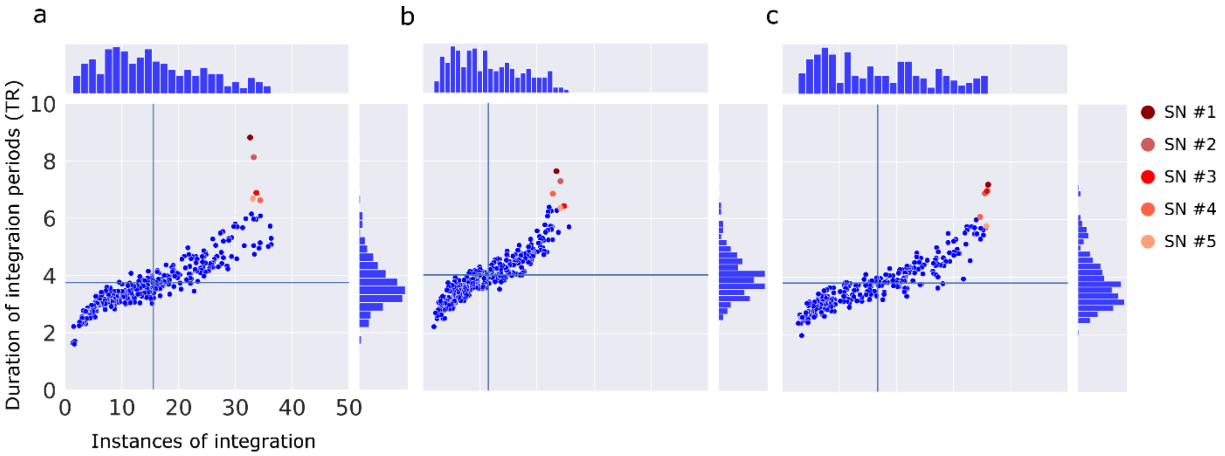


***Supplementary Figure S12.*** *Results pertaining to subnetwork (SN) integration during working memory (**a**), motor (**b**) and rest (**c**) computed on the replication "LR" data set. This figure compliments Fig. 4 in the main text.*

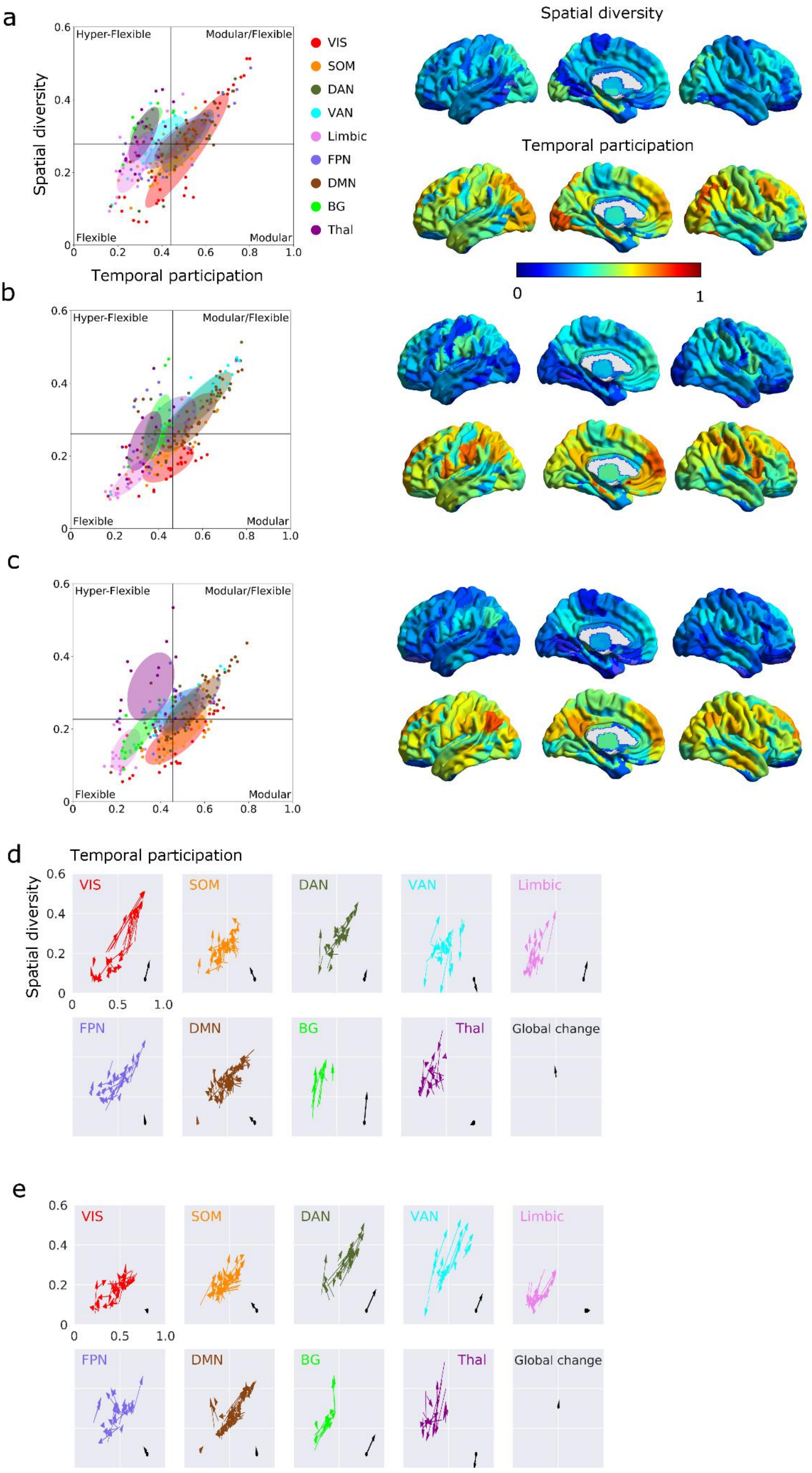
a
b
c
d
e
Spatial diversity
Temporal participation
Hyper-Flexible
Modular/Flexible
Flexible
Modular
VIS
SOM
DAN
VAN
Limbic
FPN
DMN
BG
Thal
Global change
0
1
0.6
0.4
0.2
0.8
1.0
0.5

***Supplementary Figure S13.*** *Degree of temporal participation and spatial diversity computed at the level of parcels for working memory (**a**), motor (**b**) and rest (**c**) (replication data set "LR"). This figure compliments Fig. 5 in the main text.*

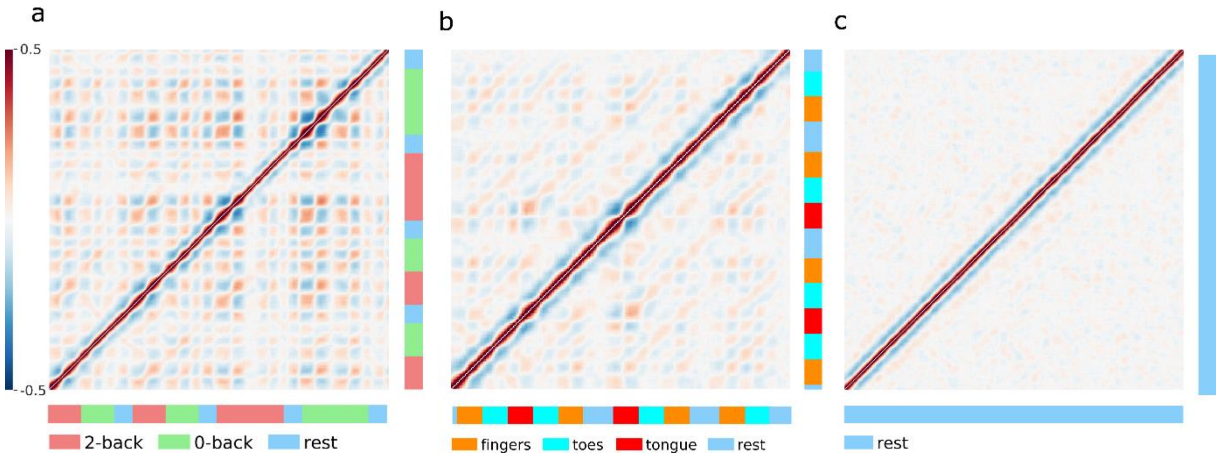


***Supplementary Figure S14.*** *Cyclicity of subnetwork integration during working memory (**a**), motor (**b**) and rest (**c**) computed on the replication data set "LR". This figure compliments Fig. 6 in the main text.*

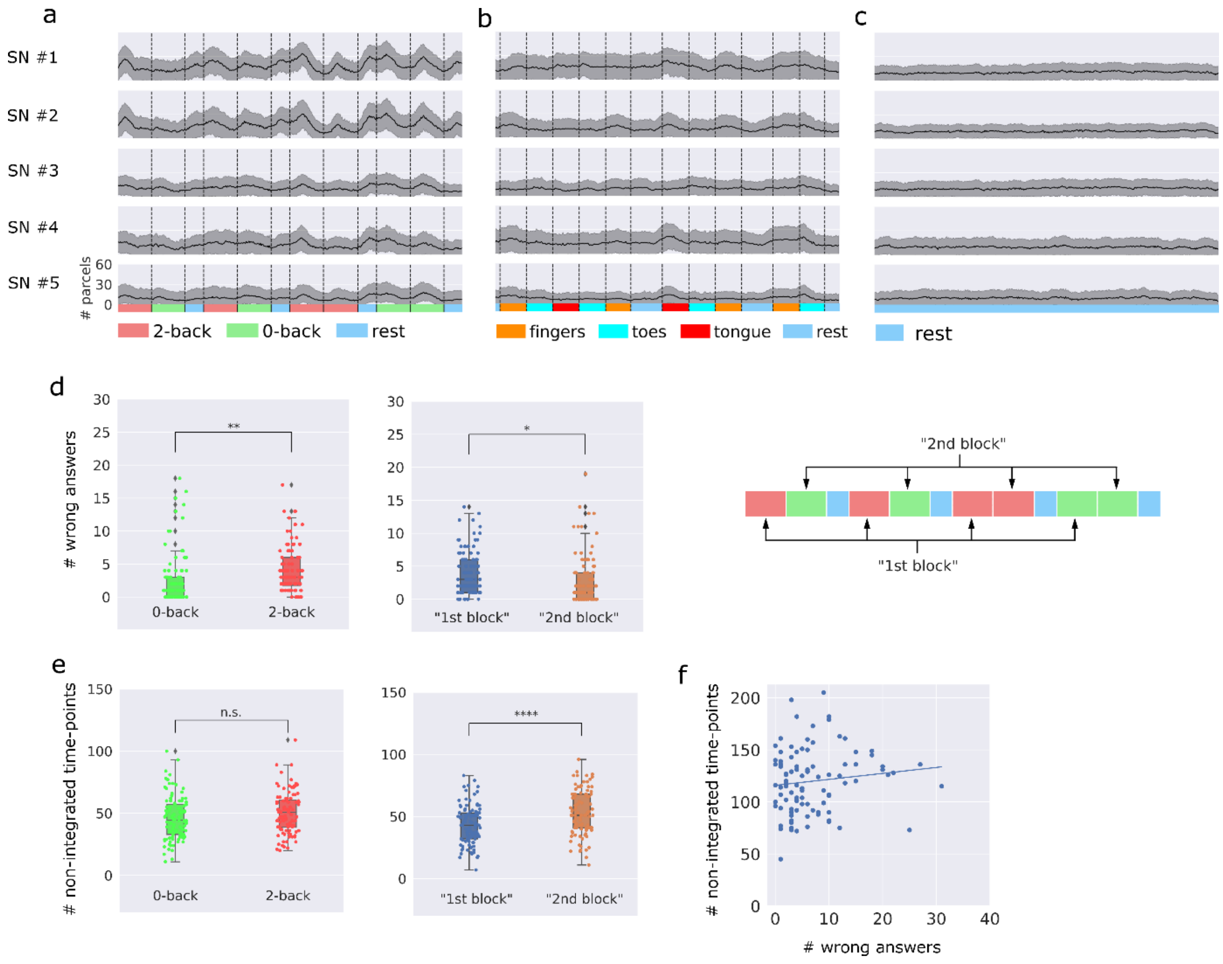


***Supplementary Figure S15.*** *Subnetwork (SN) integration and working memory task performance computed for the replication data set "LR". This figure compliments Fig. 7 in the main text. Note that the sequential order of 0-blocks and 2-blocks in this case is not balanced with respect to task difficulty, which limits the interpretability of the replication results to those presented in Fig. 7 in the main text. The correlation between the total number of non-integrated time-points for the top SN (SN #1) during the entire working memory experiment and the total number of wrong answers given by the participants were not significant (r = 0.13, p= 0.2, Spearmann correlation coefficient).*

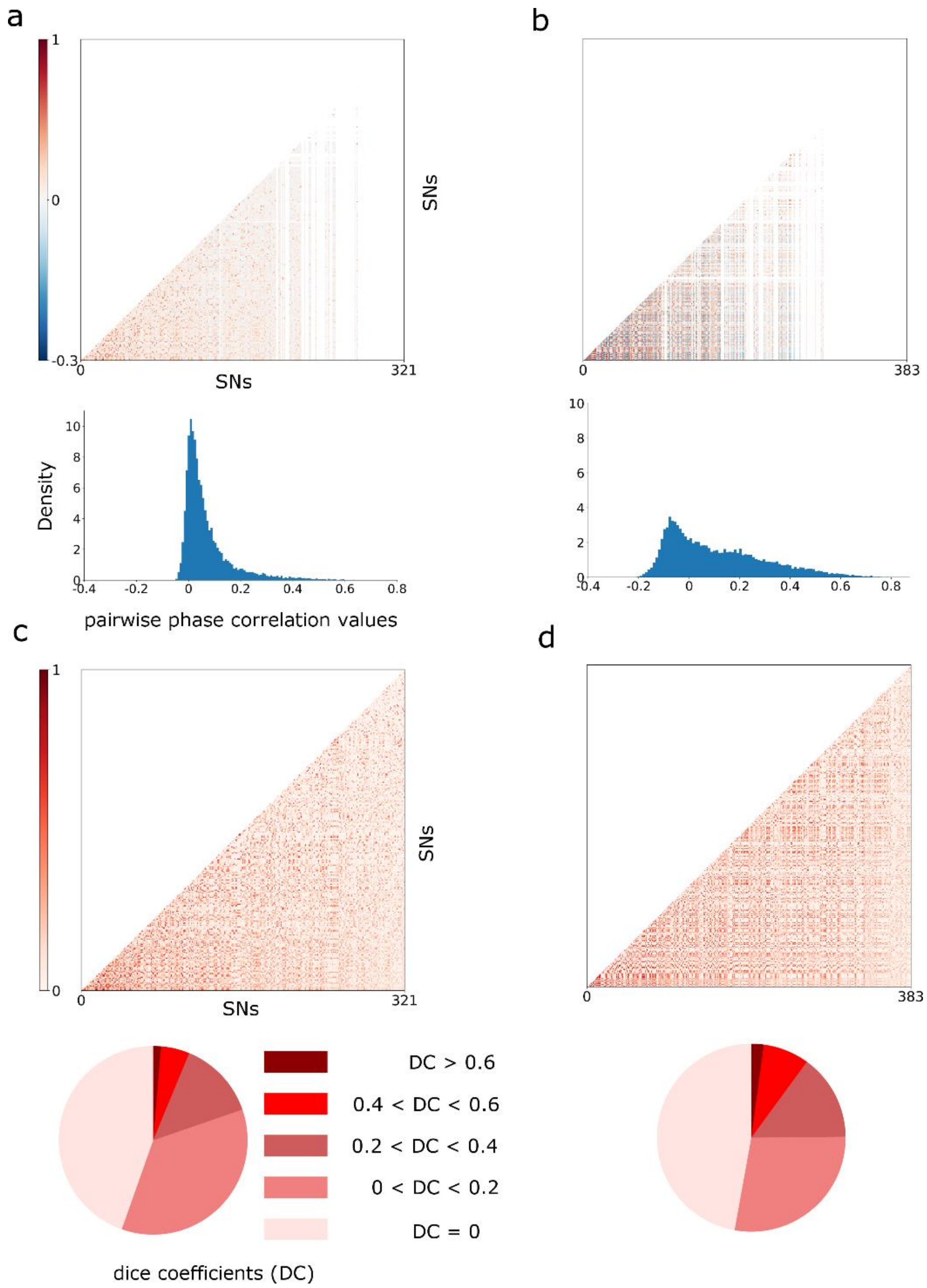


***Supplementary Figure S16.*** *Degree of phase synchronicity between all pairs of subnetworks during the working memory task here shown in the form of correlation matrices and histograms of correlation values for the faster (IMF2) (**a**) and slower (IMF4) (**b**) BOLD frequency component. Mean phase correlation values: r = 0.07 (IMF2), r = 0.11 (IMF4). The probability density is plotted on the y-axis for the histograms. The lower panel shows the degree of pair-wise spatial overlap (dice coefficient) between all subnetwork for working memory experiment for the faster (IMF2) (**c**) and slower (IMF4) (**d**) BOLD frequency components. Mean of dice coefficients: 0.10 (IMF2) and 0.12 (IMF4). For increased visibility, the relative proportion of dice coefficients (DC) were divided up into five intervals that are shown as pie charts.*

References Supplementary Material


Barch, D.M., Burgess G.C., Harms M.P., Petersen, S.E., Schlaggar B.L. et al. Function in the human connectome: Task-fMRI and individual differences in behavior. NeuroImage, 80:169-189 (2013).

Glasser, M. F. *et al.* The minimal preprocessing pipelines for the Human Connectome Project. NeuroImage 80:105–24 (2013).

Griffanti, L. *et al.* ICA-based artefact removal and accelerated fMRI acquisition for improved resting state network imaging. NeuroImage 95:232–47 (2014).

Uğurbil, K. *et al.* Pushing spatial and temporal resolution for functional and diffusion MRI in the Human Connectome Project. NeuroImage 80:80–104 (2013).

Salimi-Khorshidi, G. *et al.* Automatic denoising of functional MRI data: combining independent component analysis and hierarchical fusion of classifiers. NeuroImage 90:449–68 (2014).

Smith, S. M. *et al.* Resting-state fMRI in the Human Connectome Project. NeuroImage 80:144–68 (2013).

Van Essen, D. C. *et al.* The Human Connectome Project: a data acquisition perspective. *NeuroImage* 62:2222–31 (2012).
